# Lunar gravity predicts sleep timing

**DOI:** 10.64898/2026.01.27.701907

**Authors:** Guadalupe Rodríguez Ferrante, Leandro Casiraghi, Ignacio Spiousas, Laura Trebucq, Viridian Klei, Justin W Kahn, Alicia Rice, David Wood, Eduardo Fernández-Duque, Karen L. Bales, Cristiano M. Gallep, Diego A. Golombek, Horacio O. de la Iglesia

## Abstract

Biological rhythms are fundamental to survival, synchronizing physiological and behavioural processes like the sleep-wake cycle with predictable environmental cycles like the solar day^1^. However, the extent to which human sleep responds to the lunar cycle remains a subject of controversy, particularly given the ubiquity of artificial light and conflicting reports regarding the influence of moonlight^2–5^. While lunar illuminance has been the primary candidate for such modulation, it fails to account for semilunar rhythms^2,6^ or effects observed during the dark new moon phase^2,7^. Here, we show that human and non-human primate sleep timing is synchronized with the gravimetric cycles exerted by the Moon and Sun. By analysing longitudinal actigraphic recordings from urban cohorts in Seattle, indigenous Toba/Qom communities in Argentina, and captive titi monkeys with attenuated access to natural light, we found that sleep onset is consistently delayed around periods of maximal gravitational variations. Crucially, this synchronization aligns with gravimetric peaks regardless of whether they occur at the full or new moon. Furthermore, while human sleep timing is heavily influenced by social constraints and self-selected light exposure, this synchronization persists in titi monkeys, in which these confounds are absent. These findings identify lunar gravity as a distinct environmental cue that may regulate sleep-wake behaviour. Although this regulation may be mediated by other geophysical factors oscillating with lunisolar gravitational tides, the results suggest that the mechanism is an evolutionarily conserved trait that shapes biological timing beyond the influence of light.

## Main

Sleep is an unconscious state during which animals temporarily suspend essential behaviours for survival and reproduction, including predator avoidance, feeding, and mating. Consequently, evolutionary pressure has selected mechanisms that optimize the timing of sleep by synchronizing it to predictable environmental cycles. Indeed, the synchronization of the sleep-wake cycle with the solar day is well established across animal species and has been recognized for centuries. In contrast, the synchronization of sleep-wake cycles with the lunar cycle remains controversial, particularly in humans^5^. Recently, we showed that sleep, measured through actigraphy, in indigenous communities of the Toba/Qom people of northern Argentina, starts later and is shorter on the nights leading to the full moon, and, surprisingly, this effect is present both in communities with and without access to electric light^2^. Even more perplexing was to find an association of sleep with the lunar month in a retrospective analysis of actigraphic records of sleep from undergraduate students living in the highly urbanized and light-polluted city of Seattle^2^. These findings led us to hypothesize that the synchronization of sleep with the lunar month may not rely solely on moonlight cues but probably also on other geophysical cues associated with the lunar cycle.

Further support for this hypothesis emerged from the finding that many participants whose sleep was assessed for at least a full lunar month displayed both lunar and semilunar (∼30- and ∼15-day periods, respectively) periodicity in sleep modulation, with a delayed and shorter sleep in nights preceding the full moon, and yet again on those preceding the new moon^2^. This latter result had been previously observed in rapid-cycling bipolar patients, whose sleep and mood can change both with lunar and semilunar periodicity^6–8^.

While sleep modulation in phase with the synodic lunar month of 29.5 days could arise from people cueing on the contrasting lunar illuminance between full and new moon, sleep modulation in phase with the semilunar month cannot. Instead, what the full and new moon share is an increase in the gravitational pull (local maxima) they exert on the Earth’s surface as the Moon, Sun and Earth are aligned on the same axis, leading to semilunar spring tides.

Our central hypothesis is that the synchronization of human sleep with the lunar month is, at least partially, driven by lunar and semilunar gravity cycles. To test four predictions derived from this hypothesis, we used data from human subjects in natural settings, and from a non-human primate in a laboratory setting, as well as the corresponding gravimetric data (mean daily gravitational variation estimated based on the Moon, Sun and Earth positions). We have recorded sleep through wrist actimetry longitudinally for at least 1-3 lunar months in several cohorts of participants living in Seattle, WA, and in Toba/Qom communities from rural areas of northern Argentina during the last nine years (Table S1) and actigraphic data from titi monkeys at the California Primate National Research Centre.

### Most individuals display semilunar periodicity in sleep timing

If sleep timing is associated with lunar gravity cycles, the first prediction is that individuals will show both lunar and semilunar periodic modulation of their sleep parameters. To test this prediction, we compared the “goodness of fit” according to the Akaike criterion for each individual’s sleep parameters to a single lunar (30-day), a single semilunar (15-day), or a combined lunar and semilunar (30- + 15-day) cosinor model. The combined model with lunar and semilunar components arose as the best fit for sleep onset, offset and duration for most individuals (83-92%) of both communities (Fig. 1).

**Figure 1.**
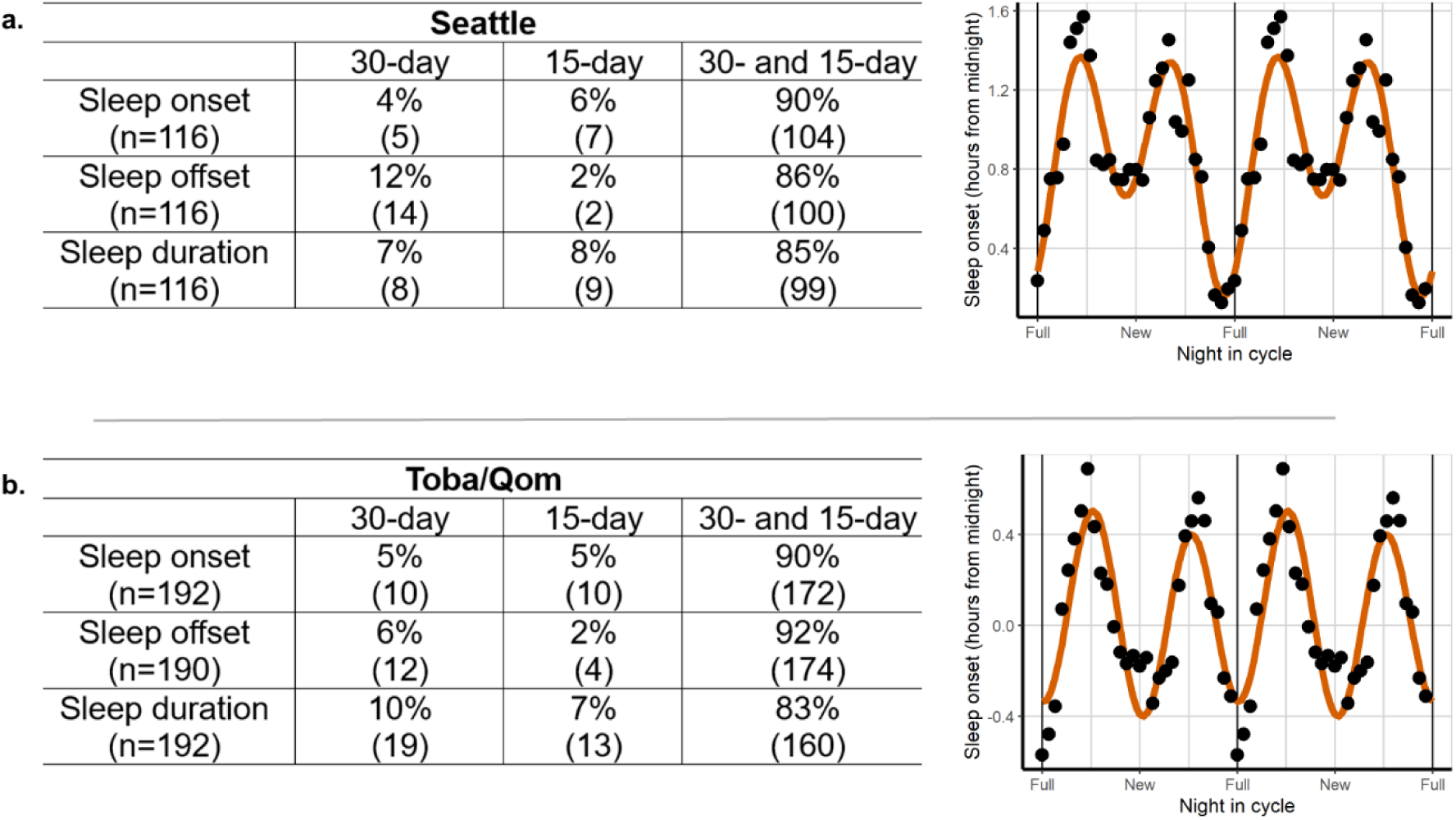
A combination of lunar and semilunar rhythmicity predicts changes in sleep timing throughout the lunar month for most individuals. The tables present the percentage of participants that better fit to one of three different cosinor models with a single 30-day period component (lunar rhythmicity), a single 15-day period component (semi-lunar rhythmicity) or including both for Seattle (a) and Toba/Qom communities (b). Plots on the right display the sleep onset of illustrative examples of individuals who present both lunar and semilunar rhythms (smoothed using a 7-day moving average and double-plotted across the moon cycle for better visualization). Note that for each sleep outcome, only participants showing at least lunar or semilunar rhythms in their sleep outcomes were included in the comparison between models. Only two Toba/Qom participants did not show said rhythms for sleep offset.

As all participants showed lunar and/or semilunar rhythmicity in their sleep onset, for the remaining of our analyses, all of them were considered.

### Delayed and short sleep can occur on the nights preceding the new moon

While the Moon’s synodic cycle determines a rhythm of illuminance tied to the full and new moon phases of the lunar month, it also drives a semilunar month of gravitational changes. Even though the variation in the gravitational pull is affected not only by the moon but also by the sun, there are local maxima or peaks during new and full moons. These peaks have different relative amplitudes, and on a given lunar month, either the full or the new moon can be associated with the highest variation in the gravitational pull (Fig. S1). If lunar gravity is associated with the timing of sleep, a second prediction is that sleep will be maximally delayed on the nights leading to the full moon within specific lunar months and maximally delayed on the nights leading to the new moon within others.

Indeed, when we pooled all our samples from each location, the Seattle cohorts showed overall delayed sleep close to the new moon (Fig. S2a; amplitude of the adjusted model was significantly different from zero [amplitude = 0.13 h, 95% confidence interval (CI) = 0.45-0.22, p = 0.003]). In contrast, the pooled cohorts from the Toba/Qom communities did not show 30-day lunar rhythmicity in their sleep onset (Fig. S2b), although there seems to be a 15-day component in this community’s sleep patterns. Results were variable for sleep offset and duration (Fig. S2a and S2b). The association between delayed sleep onset around the new moon observed in Seattle and the lack of 30-day rhythmicity in sleep onset in Toba/Qom could result from combining cohorts that present diverse phase relationships between lunar phases and their sleep onset.

To test this, we fitted a mixed cosinor model with a 30-day periodicity in each population, adding the cohort as a variable that may affect both the phase and amplitude; that is, a 30-day sinusoid was estimated for each cohort in each population (Table S2 and S3). Consistent with the prediction, cohorts showed different phases, showing later sleep onsets closer to the full moon in some cases and during the new moon in others. Strikingly, later sleep onset close to the new moon were displayed in participants in Seattle and Toba/Qom from cohorts recorded within an overlapping time window (Fig. 2). These results pointed to the need to consider each recorded cohort separately when fitting a model and to identify another variable with circalunar periodicity, other than moonlight, that could explain these changes in sleep.

**Figure 2.**
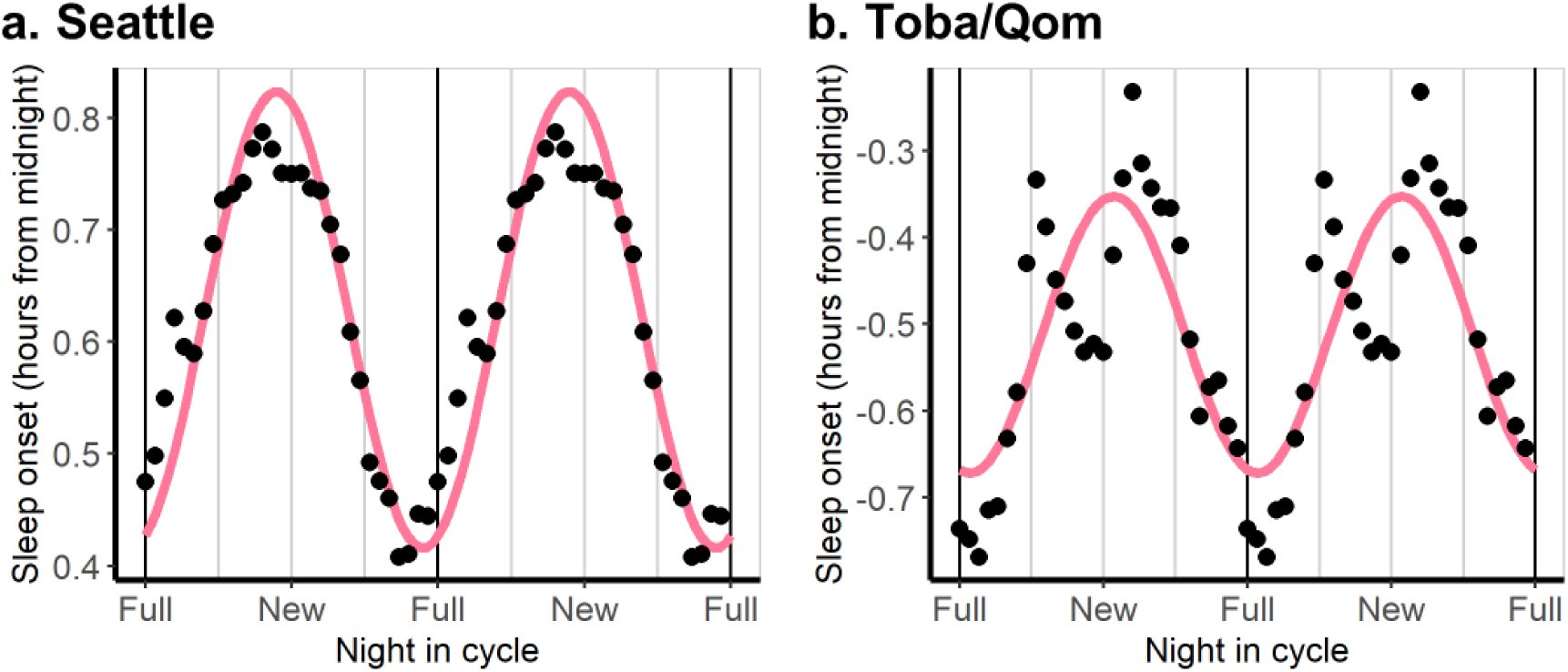
Sleep onset is later during the new moon for two overlapping cohorts in Seattle and Toba/Qom communities. In each population, we adjusted a cosinor model with a 30-day period to detect the main lunar component. We included each participant’s ID as a random factor. Purple lines show the prediction of the cosinor models. Dots represent the mean value of sleep onset for each cohort by night in the lunar synodic cycle. Data is presented as a double-plot and smoothed at the group level using a 7-day moving average for easier visualization. **A. Seattle.** Predicted bedtimes were 24.5 min later around the new moon than around the full moon (amplitude = 0.204 h, 95% CI = 0.500-0.105, p<0.0001). Data was collected from the end of February to mid-late June, 2024. N=58. **B. Toba/Qom communities.** Predicted bedtimes were 21.4 min later around the new moon than around the full moon (amplitude=0.160 h, 95% CI = 0.054-0.265, p= 0.003). Data was collected from the end of April to mid-late June, 2024. N=41.

### Local maximum of semilunar gravity predicts sleep timing

The third, and probably most important, prediction of our hypothesis is that the local gravitational variation, regardless of the phase relationship between gravity and the synodic lunar month, will predict the timing of sleep in each community and cohort. In other words, sleep parameters and local mean daily gravitational variations will show a similar waveform within each cohort and have a similar phase relationship regardless of cohort.

To study the association between the gravitational variation and sleep timing, we used a cross-correlation analysis at the cohort level. This analysis determines how well two independent oscillations linearly correlate regardless of the waveform of their oscillations. This approach overcomes the limitation of our previous cosinor analysis, which a priori assumed sinusoidal patterns for the oscillations of sleep parameters.

We first studied lunar rhythmicity of sleep parameters in three different cohorts from Seattle: Spring 2020, Winter 2021 and Spring 2024. In Spring 2024, later sleep onsets occurred one day before the new moon (acrophase = 13 night in cycle with 0 corresponding to full moon, 95% CI = 11-16), whereas they occurred 2 and 5 days before the full moon in Winter 2021 and Spring 2020, respectively (acrophase = 28 night in cycle, 95% CI = -19-15 and acrophase = 25 night in cycle, 95% CI = 21-29) (Fig. 3a and Table S2a). Notably, the absolute maximums of the mean daily gravitational variation (which combines those of the moon and the sun) are consistent with sleep onset acrophases for each cohort: they happen close to the full moon for Spring of 2020 and Winter of 2021 but close to the new moon for Spring of 2024 (Fig. 3a and 3b). Cross-correlation analysis revealed that sleep onset time and the mean daily gravitational variation at the cohort level are strongly correlated, and for all cohorts show a positive correlation without lagging any of the variables involved (i.e., lag = 0) and the maximum cross-correlation functions (CCF) were 0.51, 0.92, and 0.82 for lags of 2, 0 and -2 days for Winter 2021, Spring 2020 and Spring 2024, respectively (Fig. 3c-e). Of note, the cross-correlation is done with the mean sleep onset times for each night in cycle, not with the sinusoid predicted time (depicted for better visualization).

**Figure 3.**
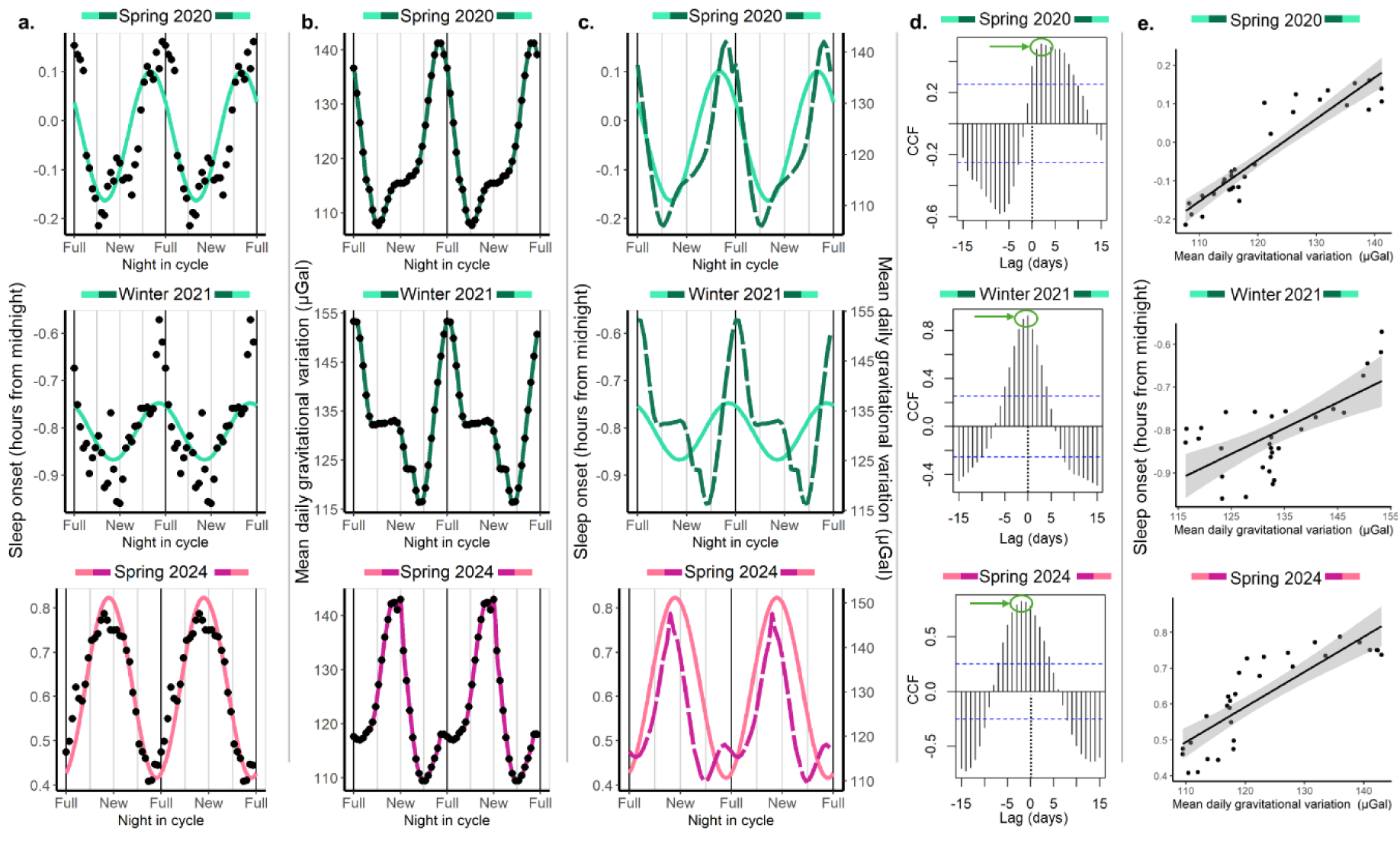
Sleep onset in Seattle shows lunar rhythmicity, and its phase depends on the cohort and is highly correlated with the mean daily gravitational variation caused by the Moon and the Sun. The colour assigned to each cohort is based on the shape of the mean daily gravitational variation and its phase relationship with the synodic lunar month. Green represents a clear maximum peak associated with the full moon while pink indicates a clear maximum peak associated with the new moon. **a. Sleep onset as a function of night in the synodic lunar cycle.** While Spring 2020 and the Winter 2021 presented later sleep onsets around/before the full moon, Spring 2024 presented later onsets around the new moon. Data are double plotted across the moon cycle for easier visualization. Dots represent the mean value by night in the cycle to which a 7-day moving average was applied. The coloured line represents the prediction of the fitted mixed cosinor model describing the relationship between night in cycle and sleep onset, where the cohort was included as a predictor of phase and amplitude. Additionally, we controlled by type of day (weekend or weekday) and we included participant id as a random factor. Note that the last panel is the same presented in Fig. 2a. **b. Mean daily gravitational variation (including Moon and Sun effects).** The absolute maxima of the mean daily gravitational variation cycle differ between cohorts, happening around the full moon for the Spring of 2020 and the Winter of 2021 and around the new moon for the Spring of 2024. Dots represent average values by night in cycle for the same dates that were included in (a). The coloured lines are for visualization, and it is a spline-based smoothening that connects consecutive dots. **c. Sleep onset and mean daily gravitational variation show similar patterns.** Coloured full lines represent the prediction of the model for sleep onset times and coloured dashed lines represent the mean daily gravitational variation. The peaks of sleep onset times either precede the peaks of gravitational variation, or they are coincident with them. **d. Cross-correlations between sleep onset and mean daily gravitational variations at the cohort level.** Plots show the cross-correlations for each cohort using the average values by night in cycle; that is, the cross-correlations between the dots represented in (a) and (b). Results are consistent between cohorts showing a positive correlation between gravity and sleep onset; that is, maximum values around a zero lag. Green circles and arrows show the obtained peaks for positive CCF values in each cohort. Peak cross-correlations lags were. Peak cross-correlations lags were: 1 for Winter 2021, -1 for Spring 2020 and -2 days for Spring 2024. CCFs above the blue doted lines statistically differ from zero. The black dotted line indicates Lag 0, that is when any signal is lagged. **e. Linear correlations between lagged sleep onset and the mean gravitational variation.** In each cohort local maxima of CCF were identified using plots in (d), the lag for which the CCF was maximum within these peaks was used for lagging sleep onset. Correlations were strong to very strong with R^2^ of 0.85, 0.44 and 0.70 for Spring 2020 (slope=0.011, 95% CI = 0.009-0.012, p<0.001), Winter 2021 (slope=0.006, 95% CI = 0.003-0.009, p=<0.001) and Spring 2024 (slope=0.010, 95% CI =0.007-0.012, p<0.001), respectively.

We obtained similar results for sleep offset, with sleep offset phases differing between cohorts (Table S2a) and later wake-up times associated with greater mean daily gravitational variation values (Fig. S3). For sleep duration, results varied according to the cohort (Table S2a). For Winter 2021, sleep onset and offset showed similar amplitudes in their lunar oscillations, while Spring 2024 sleep onset showed a higher lunar impact than sleep offset, and the opposite happened for Spring 2020. Lunar rhythmicity for sleep duration is less consistent as it is the result of sleep onset and offset timings variations: while duration changes between 14 and 18 minutes throughout the lunar month, its phase relationship with both moon phases and gravity is less consistent than for sleep onset and offset (Fig. S4).

Together, these results indicate that later sleep timings are associated with greater lunar gravity regardless of whether the maximum daily gravitational variation occurs around the full or the new moon.

We studied lunar rhythmicity of sleep onset in six different cohorts of Toba/Qom participants. Cohorts changed their sleep onset in 19-29 minutes throughout the lunar month, and these variations were statistically significant for five of them (Table S3a). Phases differ among cohorts. Winter 2016, Fall 2023 and Spring 2018 presented later sleep onsets 2 (acrophase=28, 95% CI =25-32), 8 (acrophase=22, 95% CI=18-26) and 7 (acrophase=23, 95% CI=20-25) days before the full moon. On the other hand, Spring 2023, Summer 2024 and Fall 2024 presented later onsets 7 days before (acrophase=8, 95% CI =5-10) and 2 (acrophase=17, 95% CI =11-23) and 1 (acrophase=16, 95% CI =13-19) days after the new moon, respectively (Fig. 4a and Table S3a). The cohort-specific results confirmed that the lack of rhythmicity presented in Fig. S2 was due to the cohorts oscillating in antiphase.

**Figure 4.**
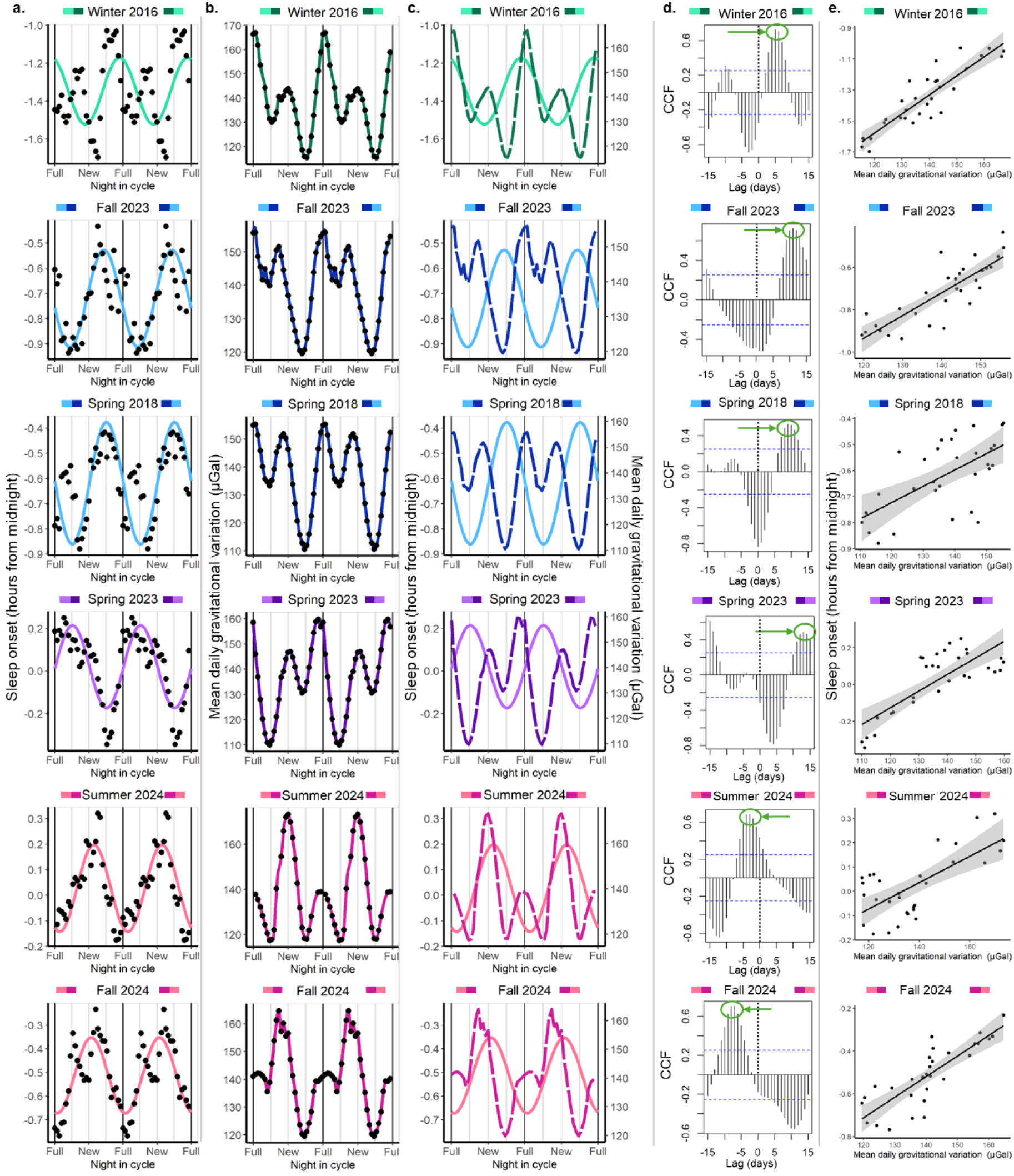
Sleep onset times in Toba/Qom communities show lunar rhythmicity, and its phase depends on the cohort and is correlated with the mean daily gravitational variation from by the Moon and the Sun. The colour assigned to each cohort is based on the shape of the mean daily gravitational variation and its phase relationship with the synodic lunar month. Green represents a clear maximum peak associated with the full moon while pink indicates a clear maximum peak associated with the new moon. When maximum peaks around the full and new moon were similar between each other, we used blue to identify cohorts with an absolute minimum before the full moon and purple cohorts with an absolute minimum before the new moon. **a. Sleep onset time as a function of night in the synodic lunar cycle.** According to the adjusted cosinor model, the only cohort whose amplitude did not differ from zero was the Summer 2024. Cohorts present different phases. Data are double-plotted across the synodic lunar month for ease of visualization. Dots represent the mean value by night in cycle to which a 7-day moving average was applied. The coloured line represents the prediction of the fitted mixed cosinor model describing the relationship between night in cycle and sleep onset, where the cohort was included as a predictor of the mesor, phase and amplitude. Additionally, we controlled by the type of day (weekend or weekday), and we included the participant id as a random factor. Note that the last panel is the same presented in Fig. 2b. **b. Mean daily gravitational variation (including Moon and Sun effects).** Dots represent average values by night in cycle for the same dates that were included in (a). The coloured lines are for visualization, and it is a spline-based smoothening that connects consecutive dots. Plots were organized according to the gravitational variation patterns, from stronger gravitational variation around the full moon to stronger gravitational variation around the new moon. **c. Sleep onset and mean daily gravitational variation show similar patterns.** Coloured full lines represent the prediction of the model for sleep onset and coloured dashed lines represent the mean daily gravitational variation. **d. Cross-correlations between sleep onset and mean daily gravitational variation at the cohort level.** Plots show the cross-correlations for each cohort using the average values by night in cycle; that is, the cross-correlations between the dots represented in (a) and (b). Green circles and arrows show the obtained peaks for positive CCF values in each cohort. Peak cross-correlations lags were: 5 for Winter 2016, 11 for Fall 2023, 9 for Spring 2018, 13 for Spring 2023, -3 for Summer 2024 and -7 for Fall 2024. CCFs above the blue doted lines statistically differ from zero. The black dotted line indicates Lag 0, that is when any signal is lagged. **e. Linear correlations between lagged sleep onset and the mean gravitational variation.** In each cohort local maxima of CCF were identified using plots in (d), the lag for which the CCF was maximum within these peaks was used for lagging sleep onset. Correlations were strong to very strong with R^2^ of 0.80, 0.71, 0.39, 0.62, 0.48 and 0.69 for Winter 2016 (slope=0.012, 95% CI =0.010-0.015, p<0.001), Winter 2023 (slope=0.011, 95% CI =0.008-0.013, p<0.001), Spring 2018 (slope=0.006, 95% CI =0.003-0.009, p<0.001), Spring 2023 (slope=0.009, 95% CI=0.006-0.012, p<0.001), Summer 2024 (slope=0.005, 95% CI=0.003-0.007, p<0.001) and Fall 2024 (slope=0.010, 95% CI=0.007-0.012, p<0.001), respectively.

Within the six Toba/Qom cohorts, three presented a clear absolute local gravity maximum around full or new moon (Winter 2016, Summer 2024 and Fall 2024; Fig 4b). In these three cohorts, sleep onset was delayed in association with higher gravitational variations, regardless of its phase relationship with the lunar synodic month (Fig. 4c-e). Interestingly, in Summer 2024 and Fall 2024, when the peak of gravity coincided with the new moon, the full moon seemed to influence the phase of the rhythm, dragging the best-fitting sleep onset sinusoid to a later phase relative to the new-moon gravitational peak (Fig. 4a, c). This may be explained by moonlight around the full moon and not by gravity.

On the other hand, and in contrast to the previous two cohorts and all the cohorts in Seattle, where gravity showed a clear absolute maximum around either the new or full moon, lunar cycles in the three remaining Toba/Qom cohorts happened to have similar gravitational peaks during the new and full moon (Fall 2023, Spring 2018 and Spring 2023). In all three cohorts, later sleep onsets happened (6-10 days) before the gravity maximum that immediately follows the absolute gravity minimum (Fig. 4c). This was reflected by negative peaks between lags 0 and 3, and positive peaks around lag 9-13 in the cross-correlations.

Results were similar between sleep offset and onset of the Toba/Qom population; that is, the latest sleep offset time still happened before or around the strongest peak of gravity in most cases. However, the phases for the two sleep parameters were different for Spring 2018 and Fall 2024. Importantly, except for the Fall 2024, the best-fitting cosinor models for sleep offset times had, overall, lower amplitudes than the ones for sleep onset times (Table S3b and Fig. S5). Finally, sleep duration results are in antiphase to sleep onset times for all cohorts, which is consistent with later sleep onsets producing shorter sleep durations (Table S3c and Fig. S6).

In summary, although the cross-correlations between sleep parameters and the mean daily gravitational variation at the cohort level demonstrated a strong association between the two variables in both Seattle and Toba/Qom communities, the phase relationships between gravity and sleep parameters are more variable between Toba-Qom cohorts than between Seattle cohorts. This may be due to the impact of a stronger semi-lunar component in both sleep and gravity and more variable patterns of mean daily gravitational variations in Toba/Qom cohorts. More importantly, the moon’s illuminance may be a more relevant cue in Toba/Qom communities, impacting the association between the gravitational variation and sleep onset.

The cross-correlation analysis performed between the mean daily gravitational variation and sleep outcomes at the cohort level has some limitations. First, sleep timing parameters represent an average of all participants, and thus, the observed cyclic pattern may not be representative of individual-level differences in the phase and amplitude of lunar patterns. Second, both the daily mean gravitational variation values and the different sleep parameters from consecutive moon cycles were averaged by day in a single, collapsed, synodic cycle, preventing the assessment of the association of gravity and sleep regardless of moon phases. Finally, as data was divided by cohort, it is difficult to conclude whether there is an overall, underlying correlation between gravity and sleep regardless of the specific lunar month of data collection. To bypass these limitations, we ran cross-correlation analysis for each participant in each cohort individually and then averaged the CCF values by lag in each population. We assessed the cross-correlations between sleep outcomes and the mean daily gravitational variation, as well as between sleep outcomes and the moon’s illuminance (Fig. 5 and Fig. S7).

**Figure 5.**
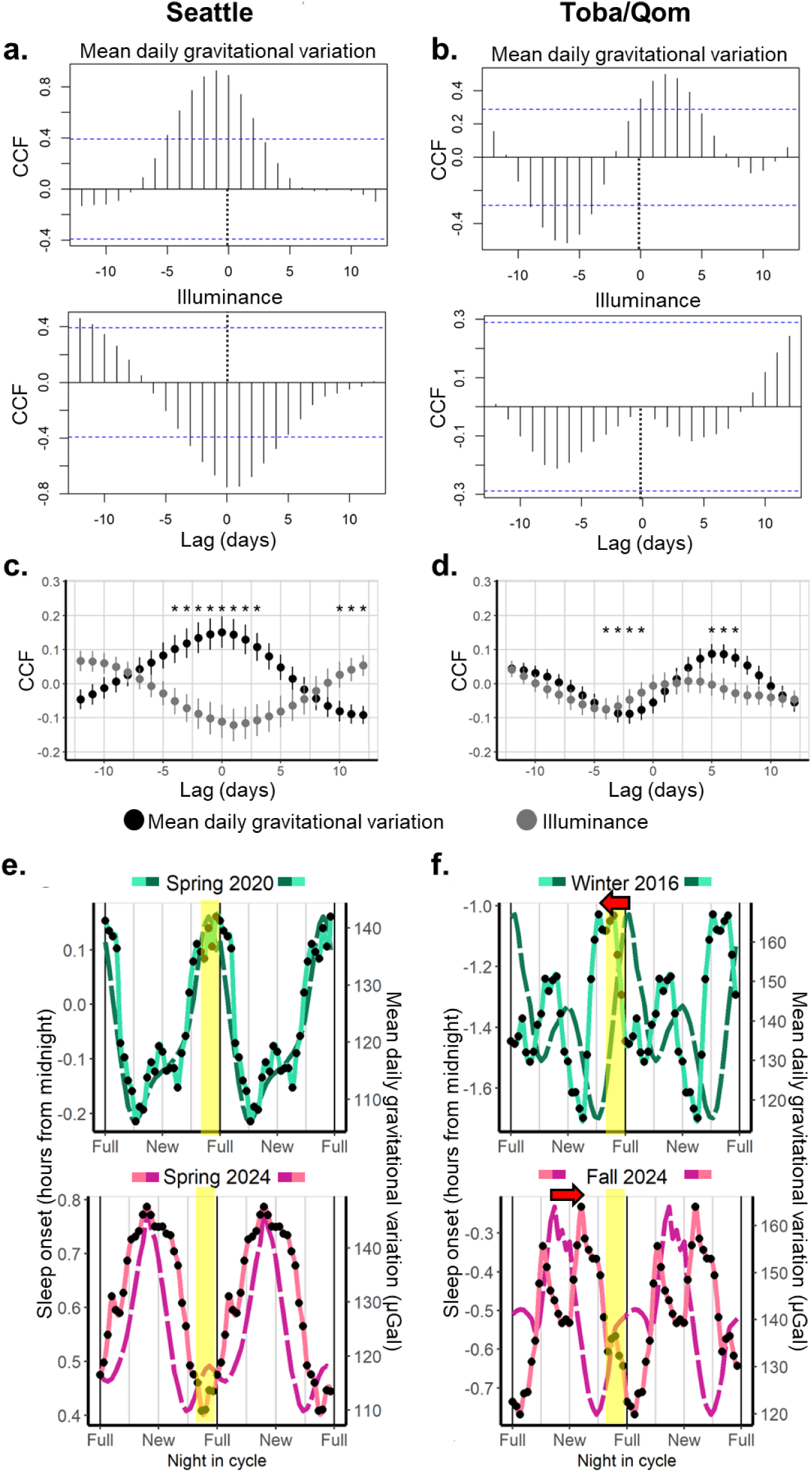
Mean daily gravitational variation but not moonlight predicts sleep onset in both communities. **a. Cross-correlation between sleep onset and mean daily gravitational variation (top) and moon’s illuminance (bottom) for one participant from Seattle. b. Cross-correlation between sleep onset and mean daily gravitational variation (top) and moon’s illuminance (bottom) for one participant form the Toba/Qom communities. c, d.** Using the results obtained at the individual level, we calculated the mean CCF for each lag and its standard error between sleep onset and both the mean daily gravitational variation (black) and the moon’s illuminance (grey) in each population. Asterisks show whether mean CCFs by lag differ from zero after controlling for multiple comparisons. **c. Seattle.** Sleep onset is positively correlated with the mean daily gravitational variation for lags around zero (-5 to 4) and negatively correlated for lags between 10 and 12. That is, sleep onset is later when the mean gravitational variation is higher. **d. Toba/Qom.** Sleep onset is positively correlated with the mean daily gravitational variation for lags 5-7 and negatively correlated for lags -4 to -1. That is, later sleep onsets precede higher mean daily gravitational variations. Lunar illuminance did not show significant correlations with sleep onset for any lag. **e, f.** Taking a cohort that presents a clear maximum around the full moon and another one that presents a clear maximum around the new moon for both Toba/Qom and Seattle, we replotted sleep onset (black dots and light coloured line) and mean daily gravitational variation (dark coloured line) as a function of the night in the synodic lunar cycle. The yellow band indicates the nights were more than 80% of the Moon is visible during the first hour after dusk. **e. Seattle.** Peaks of gravitational variation and sleep onset are centred. **f. Toba/Qom communities.** The phase relationship between sleep onset and gravitational variation is different for Winter 2016 and Fall 2024. In both cases, there is a shift (red arrows) towards nights when moonlight is available during the first hours of the night.

For Seattle, we observed that lags around zero showed positive CCFs for the mean daily gravitational variation and sleep onset; that is, sleep onsets are later around the same time that the gravitational variation is greater, consistent with what was observed at the cohort level (Fig. 5a). Sleep offset presented a similar pattern for the gravitational variation, but the maximum CCF value happened around 2-3-day lags, indicating that later sleep offsets preceded greater gravitational variations by a couple of days. Sleep durations showed lower CCF values than sleep onset and offset, and the phase was shifted but coincident with later sleep onsets and not with later wake-up times (significant CCFs around lag -4 days); that is, shorter sleep durations happened 3-5 days after high gravitational variations (Fig. S7a). There was no association between the moon’s illuminance and sleep onset (Fig. 5c), offset or duration (Fig. S7), indicating that the moon’s illuminance is not relevant to explain lunar and/or semi-lunar sleep rhythms in Seattle.

Toba/Qom participants showed positive and significant CCF values for the mean daily gravitational variation and sleep onset at lags 5-7, indicating that later onsets preceded peaks of gravitational variation, which was consistent with the results observed at the cohort level. Additionally, earlier sleep onsets happened some days after peak gravitational variations. There was no association between the Moon’s illuminance and sleep onset (Fig. 5d). Sleep offset, on the other hand, was positively associated with the moon’s illuminance but not with the mean daily gravitational variation, indicating later wake-up times some days after the full moon. Sleep duration showed a similar pattern for both the gravitational variation and the moon’s illuminance, with a 3-day shift in phase: longer sleep durations happened after the gravitational variation and the moon’s illuminance peaked, consistent with both sleep onset and offset results (Fig. S7b).

There was a difference in the phase relationship between the mean gravitational variation and sleep onset between Toba/Qom communities and Seattle; overall, CCFs between the mean daily gravitational variation and sleep onset or offset were lower for Toba/Qom communities. These results may indicate competition between moon’s illuminance and the gravitational variation in Toba/Qom. For moonlight to interfere with sleep onset, it should be bright enough and available during the beginning of the night. This happens on the nights before, but not after, the full moon. Interestingly, in Seattle regardless of when moonlight was available, mean daily gravitational variation and sleep showed a similar phase relationship (Fig. 5e). In contrast, in Toba/Qom communities the phase relationship between the gravitational variation and sleep onset seemed to differ according to moonlight availability, shifting towards the nights where moonlight was available during the first hours of the night (Fig. 5f). This may be explained by the fact that moonlight is a more relevant cue for Toba/Qom communities, which have remarkably less light pollution.

Together, our results show a clear association between sleep patterns and the mean daily gravitational variation in a highly urbanized community and a milder but still present association in a more rural context, where lunar illuminance may still impact sleep timing.

### Lunar gravity is associated with activity timing in captive titi monkeys regardless of moonlight

The fourth prediction of our hypothesis is that lunar and/or semilunar rhythms in rest-activity patterns will be associated with the lunar gravitational variation even without sociocultural zeitgebers and without self-selected exposure to electric light. We considered that non-human primates were a good model to test this prediction, given that previous field data indicated that some non-human primates, like owl monkeys, exhibit strong modulation of their circadian locomotor activity rhythms by the lunar cycle^9^, and isolating human participants in controlled environments for full months is impractical. Thus, we took advantage of the titi monkey colony at the CNPRC (Davis, California, USA) to test this prediction: we studied the lunar impact on the locomotor activity onset of captive titi monkeys that were exposed to a fixed light-dark schedule (lights on/off at 6:00 and 18:00, respectively) and had restricted access to attenuated natural light via semi-opaque skylights.

Titi monkeys showed lunar rhythmicity in their activity onset with different phases between cohorts, with later activity onsets before the full moon during Summer 2023 and before the new moon during Spring 2024 (Figure 6a and Table S4), matching the respective daily mean gravitational variation peaks (Fig. 6b). Consistently, the peak of locomotor activity onset preceded the peak of gravity in both cohorts (Figure 6c). Crucially, this synchronization occurred regardless of the availability of moonlight, as the Spring 2024 cohort delayed its activity onset close to the new moon, tracking the gravimetric peak despite the absence of lunar luminance. Together with our human data, these results show that rest-activity patterns are predicted by local gravitational variations in both human and non-human primates. Importantly, our monkey data not only rules out the possibility that lunar rhythmicity could emerge from sociocultural cues but also shows an evolutionary conservation of this rhythmicity.

**Figure 6.**
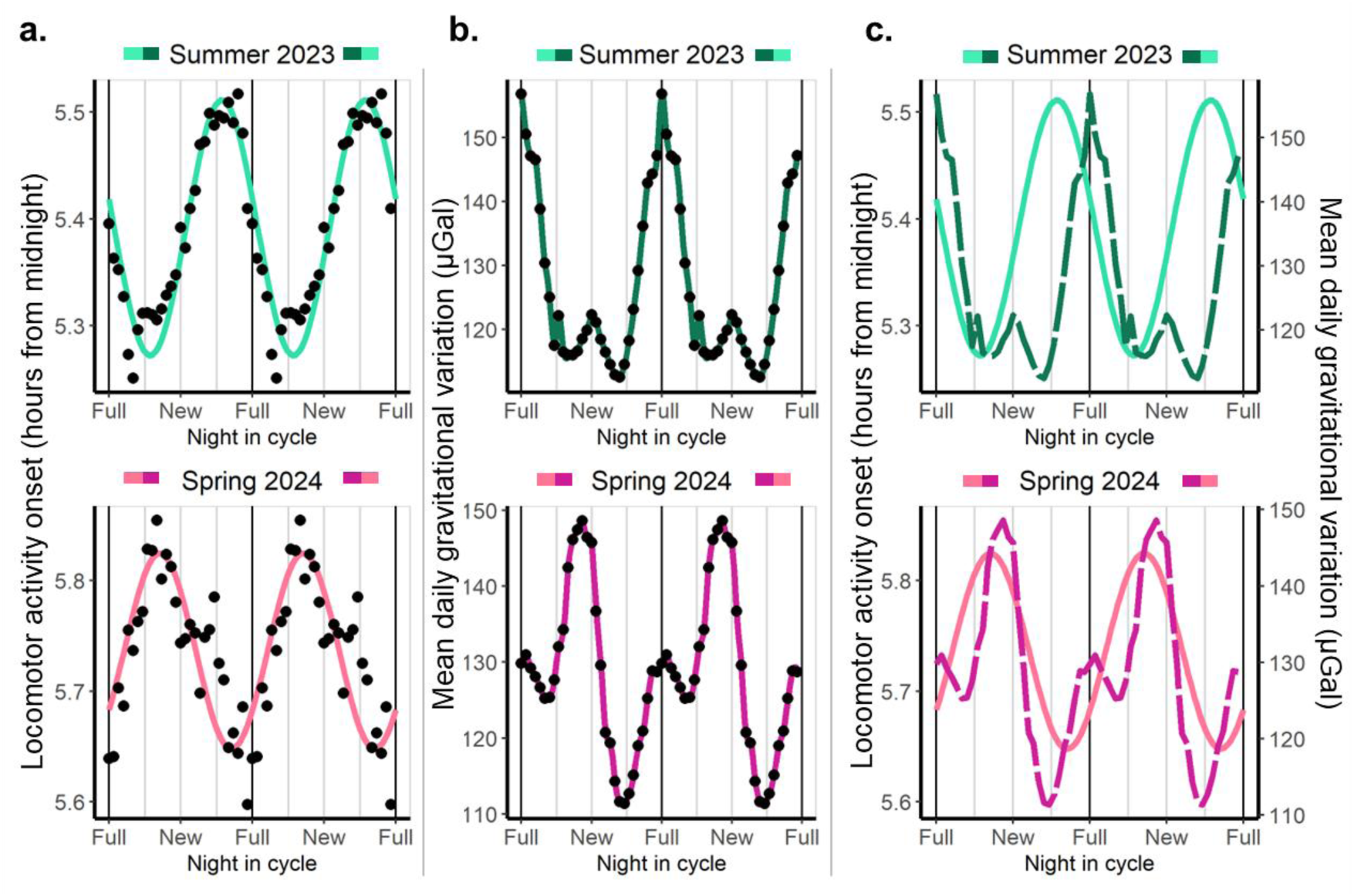
Locomotor activity onset in captive titi monkeys shows lunar rhythmicity, and its phase depends on the cohort that, coincidently, differ in the phase of the mean daily gravitational variation caused by the Moon and the Sun. The colour assigned to each cohort is based on the shape of the mean daily gravitational variation and its phase relationship with the synodic lunar month. Green represents a clear maximum peak associated with the full moon while pink indicates a clear maximum peak associated with the new moon. **a. Locomotor activity onset as a function of night in the synodic lunar cycle.** Phases differ between cohorts, with the Summer of 2023 presenting later activity onsets before the full moon and the Spring of 2024 presenting later activity onsets before the new moon. According to the adjusted cosinor model, both cohorts present an amplitude significantly different from zero. Data are double plotted. Dots represent the mean value by night in cycle to which a 7-day moving average was applied. The coloured line represents the prediction of the fitted mixed cosinor model describing the relationship between night in cycle and sleep onset, where the cohort was included as a predictor. Additionally, we controlled by the type of day (weekend or weekday), and we included the participant id as a random factor. **b. Mean daily gravitational variation (including Moon and Sun effects).** The absolute maximums of the mean daily gravitational variation differ between cohorts, happening around the full Moon for the Summer of 2023 and around the new Moon for the Spring of 2024. Dots represent average values by night in cycle for the same dates that were included in (a). The coloured lines are for visualization; they smoothly connect consecutive dots. **c. Locomotor activity onset and mean daily gravitational variation show similar patterns.** Coloured full lines represent the prediction of the model for activity onset, and coloured dotted lines represent the mean daily gravitational variation. The peak of activity onset precedes the peak of gravity, or it coincides with it.

## Discussion

Longitudinal sleep recordings showed that people living in either a highly rural or a postindustrial environment exhibit lunar rhythmicity in sleep onset, duration, and offset. These lunar monthly sleep rhythms are explained by an association between sleep parameters and the local gravitational variation exerted by both the Moon and the Sun, and in communities that rely less on electric light by an additional association with moonlight availability. Our evidence for synchronization of sleep with lunar gravity includes: (1) the presence of both lunar and semilunar rhythms in sleep measures of both communities, (2) the persistence of these rhythms in the postindustrial environment of Seattle, where moonlight is highly unlikely to serve as a cue, (3) the synchronisation, in some cohorts, of sleep parameters with new moons, with obvious minimal illuminance, that have peak monthly gravity, and (4) the presence of rest-activity lunar cycles in a non-human primate that cannot self-select light exposure and has restricted access to moonlight.

While the association between lunar gravity and sleep is present in both the Seattle and Toba/Qom communities, the CCFs were smaller in the Toba/Qom people. This attenuation may be due to at least two factors. First, during the periods in which these Toba/Qom cohorts were recorded, the mean daily gravitational variation had a stronger 15-day component than for Seattle, which leads to a lower difference between the relative maxima of gravity during the new and full moon. Second, Toba/Qom communities have very limited access to electric lighting and are likely to be more influenced by available moonlight^2,10^. When the maxima of moon’s illuminance and the gravitational variation do not coincide, these two cues may compete in shaping Toba/Qom sleep patterns. On the other hand, when the gravitational variation is similar between new and full moon, moonlight would favour the shortening of sleep around the full moon. This interpretation is supported by the finding that none of the sleep parameters were associated with moonlight in Seattle, while both sleep offset and duration were associated with moonlight in the Toba/Qom people. Indeed, our previous study in the Toba/Qom community reported later sleep onset and shorter sleep duration on the nights leading to the full moon^2^.

The data on titi monkeys, which provide a robust control for the potential influence of social cues and self-selected light exposure, further support this interpretation. In an environment in which electric light-dark schedules were fixed and access to natural light cues was restricted, the rest-activity cycles reveal high synchronization with the monthly lunar gravity cycle regardless of whether its peak occurs during the full or new moon, prioritizing gravimetric cues over moonlight.

To our knowledge, this is the first study addressing the association between sleep patterns and lunar gravity changes in healthy adults. Notably, studies in cases of rapid-cycling bipolar disorder have shown that the patients’ sleep can have high-amplitude lunar and semilunar sleep duration cycles. In turn, these sleep cycles predict mood changes, as mania and depression are associated with short and long sleep duration, respectively^6,8^.

Evidence in women points to a relationship between human physiology and lunar gravity. The menstrual cycle can be synchronized with the lunar month, and this synchronization involves not only the synodic lunar phases but also two lunar cycles that are primarily gravimetric, the anomalistic and tropical lunar months^11,12^. Moreover, similarly to what we observed in Seattleites’ sleep, menstrual synchrony with the full moon weakens in women living in urban areas, although they still show synchrony in lunar months with high gravimetric amplitudes^12^. Interestingly, the evidence from both bipolar patients and women’s menstrual cycle points to the existence of an endogenous timing system weakly entrained to lunar gravity^7,11–13^. This is consistent with our observation that most individuals present a lunar rhythm in their sleep, but variability in the phase relationship with the lunar cycle is high. However, the unequivocal demonstration of an endogenous circalunar clock would require months-long isolation of human subjects from lunar light and gravity cycles, a virtually impossible task.

The synchrony between human sleep-wake patterns and the synodic month has remained controversial, as several studies reported its absence^3,5,14,15^. Together with the lack of longitudinal recordings^2,16^, this may reflect that prior studies did not account for gravitational cycles. As shown in this study, combining data from cohorts that differ in their gravitational patterns can make the association between sleep and the lunar month disappear. Interestingly, most of the studies that have found an association between human sleep and moon phases reported that full moon is associated with later sleep onset, longer sleep latency and shorter sleep duration^2,4,17–20^ This may indicate an impact of moon phases on sleep independent of gravity, as moonlight is not the only variable that differs between moon phases (e.g., the effect of the Moon on the Earth’s magnetosphere differs between new and full moon^21^). Nevertheless, given that we showed here that some cohorts exhibited the opposite phase relationship, and a recent paper reported earlier bedtimes around the full moon for one of their study groups^17^, this may also reflect a publication bias, especially because the prevailing hypothesis is based solely on an effect of moonlight.

The mechanisms by which humans could detect gravitational cycles remain unknown and enigmatic, particularly considering that the perceived gravitational variation is dominated by the effect of the Earth while the magnitude of the gravitational variations caused by the Moon and the Sun are relatively small. However, the Moon and the Sun exert highly predictable, cyclic variations in gravity to which all organisms have been exposed since the genesis of life, which makes them relevant as potential rhythmic environmental cues. Moreover, although here we show a correlation between gravitational cycles and sleep, an alternative synchronization mechanism could involve not gravity itself but other geophysical factors covarying with gravity. Lunar and solar gravitational cycles are associated with ocean, solid earth, and atmospheric tides, as well as with fluctuations in geomagnetic and electric fields^22–29^. It has long been known that several living organisms are able to sense and respond to these types of environmental cues. For example, birds and turtles use magnetic fields for orientation, and two main detection mechanisms have been proposed: retinal cryptochromes and iron-rich particles (e.g., magnetite)^30–32^. Evidence in mammals has been more elusive, but recent findings show that mouse retinal cryptochrome responds to magnetic fields and mediates behavioural changes in response to such fields^33^. This raises the possibility that humans, too, may be sensitive to subtle geophysical cues, even if the underlying sensory pathways remain poorly understood.

Humans have likely benefited from being active during moonlit nights since ancestral times, either to be able to avoid predators or to exploit extended foraging or social opportunities^34,35^. In contrast, there is no clear adaptive advantage for sleep to be directly modulated by gravitational variations. One possibility is that stronger gravitational tides increase sensitivity to nocturnal light, which could, in turn, delay activity offset when light is available at night. Under natural light conditions, this mechanism would predict inhibition of sleep onset around both new and full moons, while increased nocturnal activity would be expected primarily during moonlit evenings. Intriguingly, our data on titi monkeys and previous results in nocturnal monkeys^9^ suggest that the modulation of sleep by the lunar month may represent an evolutionarily conserved trait in primates.

This study has several limitations that should be acknowledged. First, it is observational in nature and based on statistical correlations, which prevents us from drawing causal inferences about the relationship between gravitational cycles and sleep. Second, neither the gravitational values used in our analysis nor the predicted moonlight levels were directly measured but estimated from the relative positions of the Moon, the Sun, and the Earth. While these estimates are well established, they may not fully capture local fluctuations in geophysical variables (e.g., the impact of bodies of water on local gravity and the cloudiness on moonlight). Third, although our dataset includes longitudinal recordings, the maximum sample duration was ten weeks. Longer-term data would allow us to examine whether the synchronization between sleep, gravity, and moonlight changes within the same individual over extended periods of time.

Despite these limitations, this study has notable strengths. First, by using estimated daily numerical values of daily gravitational variation, we advance beyond previous approaches that relied solely on lunar monthly phases or periodicities, thereby providing a more rigorous and quantitative assessment of gravitational influences. Second, we include three distinct populations that differ socioculturally, in their access to electric light and in their exposure to moonlight (Seattle and Toba/Qom participants and captive titi monkeys), which allows us to test the generalizability of our findings across diverse ecological contexts. Finally, the integration of both human and non-human primate data underscores the potential evolutionary relevance of gravitational cues for sleep regulation.

In this work, we present evidence for a novel association between human and non-human primates’ rest-activity patterns and the gravitational variation exerted by the Moon and the Sun, raising new questions about its physiological relevance. Future studies are required to determine whether these cycles influence additional aspects of human physiology and health, and to elucidate the mechanisms through which such effects may occur. Since humans are continuously exposed to predictable variations in gravity, understanding how these forces interact with biological systems could have broad implications, potentially opening the way to the development of novel diagnostic or therapeutic strategies.

## Methods

### Participants and study groups

For human participants, all study procedures were approved by the Internal Review Board of University of Washington’s Human Subjects Division and were in agreement with the Declaration of Helsinki. Written consent was obtained from participants of Seattle, all of which were 18 years old or older. Oral consent was obtained from Toba/Qom participants after a verbal explanation of all procedures in Spanish with the help of a Toba/Qom translator whenever necessary. Parental oral consent was obtained for participants under 18 years old, who also gave their assent to participate.

Seattle participants were unaware of any relationship between their sleep study and the moon. Toba/Qom participants were aware that we were interested in studying the relationship between the moon cycle and sleep but were unaware of any of our specific predictions. Moreover, they were unaware of our interest in the association between sleep patterns and changes in the gravitational pull.

For the non-human primate subjects, we received all necessary approvals from the Institutional Animal Care and Use Committee of Yale University, the University of California, Davis, and the California National Primate Research Center for this investigation.

### Seattle participants

Participants from Seattle were 100 adults (63 females, 6 unreported, mean age: 28.5, range: 18.5-74.9; 5 participants did not report their age), who wore one of two models of wrist actimeter (Actiwatch Spectrum Plus, Respironics, OR, USA; or AX3, Axivity, Newcastle, UK) for 4 to 10 weeks. Only those who had provided valid actigraphic data for more than 24 days (more than 80% of the synodic lunar cycle) were included. Additionally, nights with sleep durations above or below 2.5 median absolute deviations of the sample were excluded. The mean (± SD) number of nights per participant was 40.9 (±15.7). Participants also completed an online sleep diary. Data were collected for five different cohorts with a few participants being included in more than one (Table S1a). Only cohorts with 10 participants or more were included in the analyses.

### Toba/Qom participants

Participants were 101 individuals (59 females, mean age: 23.2, range: 12.1 to 50.9) from indigenous Toba/Qom communities in Formosa, Northern Argentina. Data were collected for seven different cohorts, with some participants being in more than one cohort (Table S1b). Only cohorts with 10 participants or more were included in the analyses.

Toba/Qom participants belonged to three different communities, one settled in the outskirts of Ingeniero Juárez (pop. ∼13,000), and two settled in isolated rural areas 60 km from the former (see Casiraghi et al. 2021 for details^2^). These communities live under very limited socioeconomic conditions and spend much of the daytime outdoors and exposed to natural daylight. While originally only the Ingeniero Juárez community had access to electricity, light has been introduced in rural areas starting in 2016, and currently all communities have electric power at their homes^36^. Participants wore a wrist actimeter (Actiwatch Spectrum Plus, Respironics, OR, USA; or AX3, Axivity, Newcastle, UK) on their non-dominant arm and completed a sleep diary for 1 to 2 months at a time (40.9 days ± 9.6). In each cohort, only those participants who provided valid actigraphic data for more than 24 days (more than 80% of the lunar cycle) were included. Additionally, diurnal sleep events (before sunset), which have low frequency (0.01% of the total sleep events), were excluded and nights with sleep durations above or below 2.5 median absolute deviations were excluded.

### Titi Monkeys

The animals were 20 (10 females) titi monkeys (*Plecturocebus sp.*) living in captivity under a 12-12 light-dark cycle regime (lights were on from 6am to 6pm) in the California National Primate Research Center, Davis, California. All animals were fitted with triaxial AX3 accelerometers attached to ball chain collars (3mm diameter) using a food-safe, electrician’s shrink polyolefin tube. Monkeys wore them for 3-4 weeks at a time, and data were collected while they wore it or until the devices ran out of battery. Animals averaged 50.35 (±29.03) days of tracking, however including eventual gaps in recordings due to devices running out of battery, thus the data are split into shorter recordings of around 25 days each. Data included was collected between June 10 and August 31, 2023 (n=9) and February 14 and June 30, 2024 (n=11). Animals were housed under a fixed 12:12 light-dark cycle (lights on at 06:00, lights off at 18:00). While the primary photoperiod was artificially controlled, the facility contains semi-opaque skylights that permit the entrance of attenuated natural light.

### Data management and analyses

All the data management and analyses were performed in the R statistical coding language (version 4.4.1)^37^.

### Activity and sleep outcomes

For Actiwatch loggers, data acquisition interval was set to 1 min. Recorded data were downloaded and exported using the Philips Respironics Actiware software V.6.0.9. The Actiware software was set to determine sleep onsets whenever 10 consecutive minute bins were classified as of immobility (<4 activity counts). Sleep offsets are the first minute bin classified as “wake” (> 40 activity counts) after the last 10 consecutive minutes of immobility in a sleep bout.

For AX3 loggers, data acquisition frequency was set in 50Hz and a range of 8 ng. Data was saved as tri-axial accelerometer data in g.

For human participant data, raw data were processed using ‘GGIR’ package (version 3.1.2) for R^38^. Sleep metrics were derived using GGIR’s HDCZA (Heuristic algorithm looking at Distribution of Change in Z-Angle) algorithm based on sustained inactivity. Nights were included if they contained at least 16 hours of valid data within a 24-hour noon-to-noon window. Sleep onset and offset times, as well as total sleep duration, were extracted from the GGIR output for downstream analysis.

Sleep diary responses were used for the validation of the calculated sleep outcomes and to discard data from time ranges when the subjects were under different conditions to those in their main study group (e.g., traveling).

For non-human primates, raw data were used to calculate the Vectorial Dynamic Body Acceleration (VeDBA), a highly used and validated proxy for animal locomotor activity^39,40^. We used this proxy to estimate each animal’s daily activity onset. For that, we evaluated all possible step functions representing a transition from rest to activity (defined as a change from 0 to 1 at a candidate onset time) and identify the one that shows the highest correlation with the binarized activity signal (where values below the daily mean are coded as 0 and those above as 1). To check the performance of our estimation, we built actograms and visually checked that the calculated values were consistent with a visual estimation of the activity onset.

### Lunar phases and night in cycle

For each day, we obtain the percentage of the illuminance of the moon using the ‘suncalc’ package in R^41^.

We assign values from 0 to 29 to each night according to the position of that night in the lunar synodic cycle (lunar phases cycle). We used the percentage of the illuminance of the moon to do this assignment. A completely, or almost completely, illuminated moon corresponds to a full moon and was assigned the number 0 for night in cycle, while a (almost) completely not illuminated moon corresponds to a new moon and was assigned the number 14 for night in cycle.

### Gravimetric tide predictions

Predictions of the local gravimetric tide (dg) were obtained for each study location using the executable software D-Tide (see Appendix at the Repositório de Dados de Pesquisa da Unicamp, https://doi.org/10.25824/redu/UGMCJV42), which is based on the theoretical model developed by Longman^43^. Using the orbital mechanics of the Sun–Earth–Moon system, the model estimates temporal variations in the gravitational acceleration experienced at Earth’s surface due to lunisolar forcing, computed separately for the horizontal and vertical components of the gravimetric tide at any geographic location. The vectorial sum of these components represents the total lunisolar gravimetric tide (dg), expressed as a deviation from Earth’s mean gravitational acceleration (∼9.8 m/s²). Importantly, dg reflects small but predictable cyclic departures from Earth’s constant gravitational field arising from the relative positions of the Moon and Sun, rather than changes in Earth’s baseline gravity itself.

The model requires as input a start date, temporal resolution (hours), duration (days), and geographic coordinates. Hour-by-hour estimates of dg were generated and subsequently averaged to obtain a mean daily value of the total lunisolar gravimetric tide. Throughout the manuscript, this quantity is referred to as the mean daily gravitational variation. Because the main cause for the lunar and semilunar components (peaks around new and full moons) in the changes in lunisolar gravity is the monthly lunar movement, we also generically refer to the lunisolar gravity as lunar gravity.

### Statistical analysis

An important part of the analysis was done using cosinor models. Cosinor analysis is a regression-based approach commonly employed to quantify periodic signals in biological time series. In this framework, the observed data are modeled as a sinusoidal function with a predefined period, allowing estimation of three interpretable parameters: the mesor (rhythm-adjusted mean), the amplitude (half the peak-to-trough difference, reflecting rhythm strength), and the acrophase (timing of the peak relative to the cycle). Cosinor models are particularly well suited for detecting and comparing cyclic variation when the period of interest is known a priori, as is the case for environmental rhythms such as daily or lunar cycles.

At the individual level, we examined the association between sleep habits (onset, offset, and duration) and the lunar synodic cycle in each population (Seattle and Toba/Qom) by fitting cosinor models separately to each participant. To assess whether sleep habits exhibited lunar (30-day), semi-lunar (15-day), or combined (30- and 15-day) rhythmic patterns, we fitted three alternative models to each participant’s data: one including a single sinusoid with a 30-day period, one including a single sinusoid with a 15-day period, and one including the sum of two sinusoids with 30- and 15-day periods. In the models including two sinusoidal components they were not phase-locked, that is, the phase relationship between the two components was determine by the best fitting model. Prior to modeling, data were averaged by day in the lunar cycle and smoothed using a 7-day moving average window to reduce high-frequency noise.

Among participants exhibiting a significant association between the lunar cycle and their sleep habits (defined as an amplitude statistically different from zero in at least one of the fitted cosinor models) we compared the three models using the Akaike Information Criterion (AIC). Participants were classified according to the model providing the best fit, and we report the proportion best described by lunar, semi-lunar, or combined rhythmic patterns, with particular emphasis on whether sleep habits were more accurately captured by a combined 30 + 15-day model relative to models including a single periodic component. To assess the robustness of this approach, we repeated the individual-level analyses using shorter smoothing windows (3 and 5 days). While the proportion of participants best fit by each model varied with window size, reflecting differences in noise reduction, the overall qualitative pattern was preserved, with evidence for both lunar (30-day) and semi-lunar (15-day) components at the individual level across windows. Accordingly, the main conclusions regarding the presence of lunar and semi-lunar modulation were not dependent on the specific choice of smoothing window.

At the whole sample level, we examined the association between sleep habits (onset, offset, and duration) and the lunar synodic cycle by fitting mixed cosinor models with a 30-day period. For each model, a sleep parameter was included as the dependent variable and the night in the synodic cycle as the predictor. We included participant id as a random factor. Additionally, we controlled by type of day (weekday or weekend), sex and age. When studying this association for each cohort, we included the cohort as the group in the model, meaning that one sinusoid was fitted to each cohort. In the case of titi monkeys, we fitted a similar but simpler mixed cosinor model: locomotor activity onset was included as the dependent variable, the night in the synodic cycle as the predictor, cohort was included as the group and subject id was included as a random factor.

For all cosinor analysis, we used the GLMMcosinor package (version 0.2.0) in R^44^. We plot the prediction of the models using colored lines on top of the mean values for each sleep outcome by day in the synodic cycle.

For human participants and given that the phase of the sleep rhythms differs between cohorts, to study whether these rhythms throughout the lunar synodic month are associated with gravitational variations, we run a cross-correlation at the cohort level between the mean values of sleep outcomes (sleep onset, offset and duration) and the mean daily gravitational variation by night in cycle of the synodic month.

Finally, for the Seattle and Toba/Qom populations, we computed cross-correlations between sleep outcomes and both lunar illuminance and mean daily gravitational variation at the individual level, separately within each population. For each lag, cross-correlation functions (CCFs) were averaged across individuals and compared against zero, with multiple comparisons controlled using the Benjamini-Hochberg procedure.

Prior to computing CCFs, up to three consecutive missing daily values were imputed using linear interpolation in time. Only recordings containing at least 30 consecutive days of data were included in the analyses. To attenuate high-frequency day-to-day variability and account for weekly structure in sleep timing, sleep recordings were smoothed using a 7-day moving average. To assess the robustness of the results to the choice of smoothing window, all analyses were repeated using a 3-day moving average, yielding comparable results. Additionally, to evaluate whether the observed associations could arise spuriously from shared temporal structure, we conducted a permutation analysis in which sleep onset values were randomly shuffled within each individual while preserving the original temporal structure of the lunar and gravitational signals. No statistically significant cross-correlations were observed in the shuffled data.

## Acknowledgments

We are thankful to A. García, M. Pérez, K. Ortiz, M. Rotundo, E. R. Guiñazú, and N. West for field assistance and the Toba/Qom people for the tremendous support and collaboration. <u>Funding:</u> This study was supported by NSF RAPID award #1743364 to E.F.-D. and H.O.d.l.I.; by Leakey Foundation grant 1266 to H.O.d.l.I.; CNPRC base grant (NIH P51OD011107) and NIH R01HL162311 to H.O.d.l.I.

## Author contributions

G.R.F. designed the study, collected Seattle data, analysed data, and wrote the manuscript. L.C., I.S. and L.T. collected Toba/Qom data, organized and processed Toba/Qom recordings, and revised the manuscript. V.K., J.W.K. and A.R. collected, organized and processed Seattle recordings. D.W. collected titi monkey data. E.F.-D. and K.L.B. provided resources for the monkey studies and revised the manuscript. C.M.G. provided the gravitational data and revised the manuscript. D.A.G. provided resources and revised the manuscript. H.O.d.l.I designed the study, provided resources, collected Toba/Qom and Seattle data, and wrote the manuscript.

