## Supplementary material for "Lunar gravity predicts sleep timing"

### Supplementary Tables

**Supplementary Table 1. Studied cohorts' information.** The table shows the number of participants, the date that data collection started and finished and the mean number of nights per participant for each cohort for each population under study. **a. Seattle.** Five cohorts were studied and two. **b. Toba/ Qom communities.** Seven cohorts were studied.

**a. Seattle**

| **Cohort** | N | Start date | End date | Mean numbers of nights |
|---|---|---|---|---|
| Spring 2020 | 32 | 2020-03-28 | 2020-06-10 | 38 |
| Winter 2021 | 13 | 2021-02-15 | 2021-03-16 | 28 |
| Spring 2023 | 7 | 2023-03-29 | 2023-05-22 | 53 |
| Spring 2024 | 58 | 2024-02-29 | 2024-06-19 | 45 |
| Fall 2024 – Winter 2025 | 6 | 2024-10-01 | 2025-01-08 | 27 |

**b. Toba/Qom**

| **Cohort** | N | Start date | End date | Mean numbers of nights |
|---|---|---|---|---|
| Winter 2016 | 32 | 2016-08-29 | 2016-09-30 | 31 |
| Spring 2017 | 9 | 2017-10-01 | 2017-11-06 | 33 |
| Spring 2018 | 25 | 2018-10-16 | 2018-12-13 | 48 |
| Fall 2023 | 20 | 2023-05-08 | 2023-06-29 | 44 |
| Spring 2023 | 45 | 2023-10-12 | 2024-12-08 | 46 |
| Summer 2024 | 17 | 2024-01-24 | 2024-02-20 | 27 |
| Fall 2024 | 44 | 2024-04-30 | 2024-06-20 | 44 |

**Supplementary Table 2. Summary of the cosinor models testing lunar rhythmicity in Seattle's cohorts sleep patterns.** The models include only one sinusoidal component with a period of 30 days. In each, one sleep outcome (sleep onset, offset, or duration) was included as the dependent variable, while night order in the lunar synodic cycle was the time variable and the cohort as the group. Additionally, we control for type of day (weekday/weekend), cohort, sex and age, including them as modifiers of the mesor. Winter 2021 cohort and weekdays are the reference levels. Significant amplitudes are shown in bold font. The intercept, amplitudes and the effects of sex, age and type of day are reported in hours for sleep onset and offset and in minutes for sleep duration. Acrophases are reported in radians. **a. Sleep onset.** Only the Spring 2024 cohort showed a significant amplitude. Sleep onsets were represented as hours from midnight. **b. Sleep offset.** Only Spring 2020 cohort showed a significant amplitude. Sleep offsets were represented as hours from midnight. **c. Sleep duration.** Both Spring 2024 and Spring 2020 cohorts showed a significant amplitude. Sleep duration was measured in minutes.

| **a. Sleep onset** | | | | | |
|---|---|---|---|---|---|
| | Estimate | SE | Lower CI | Upper CI | p |
| Intercept | 1.383 | 0.521 | 0.362 | 2.404 | 0.008 |
| Type of day (weekend) | 0.387 | 0.046 | 0.297 | 0.477 | <0.001 |
| Sex | 0.718 | 0.262 | 0.205 | 1.232 | 0.006 |
| Age | -0.054 | 0.013 | -0.080 | -0.029 | <0.001 |
| Winter 2021 | -0.363 | 0.092 | -0.543 | -0.184 | <0.001 |
| Spring 2024 | 0.079 | 0.315 | -0.538 | 0.697 | 0.802 |
| Amplitude – Spring 2020 | 0.132 | 0.077 | -0.019 | 0.283 | 0.086 |
| Amplitude – Winter 2021 | 0.060 | 0.118 | -0.172 | 0.291 | 0.614 |
| **Amplitude – Spring 2024** | **0.204** | **0.052** | **0.102** | **0.306** | **<0.001** |
| Acrophase – Spring 2020 | -1.027 | 0.458 | -1.924 | -0.129 | 0.025 |
| Acrophase – Winter 2021 | -0.456 | 1.796 | -3.976 | 3.063 | 0.799 |
| Acrophase – Spring 2024 | 2.812 | 0.226 | 2.368 | 3.256 | <0.001 |
| **b. Sleep offset** | | | | | |
| | Estimate | SE | Lower CI | Upper CI | p |
| Intercept | 8.718 | 0.484 | 7.769 | 9.667 | <0.001 |
| Type of day (weekend) | 0.811 | 0.048 | 0.718 | 0.905 | <0.001 |
| Sex | 0.222 | 0.246 | -0.261 | 0.704 | 0.369 |
| Age | -0.031 | 0.012 | -0.055 | -0.007 | 0.010 |
| Winter 2021 | -0.223 | 0.095 | -0.409 | -0.036 | 0.019 |
| Spring 2024 | -0.209 | 0.294 | -0.786 | 0.367 | 0.477 |
| **Amplitude – Spring 2020** | **0.168** | **0.065** | **0.040** | **0.295** | **0.010** |
| Amplitude – Winter 2021 | 0.143 | 0.114 | -0.080 | 0.367 | 0.208 |
| Amplitude – Spring 2024 | 0.110 | 0.057 | -0.001 | 0.221 | 0.051 |
| Acrophase – Spring 2020 | -1.892 | 0.478 | -2.829 | -0.956 | <0.001 |
| Acrophase – Winter 2021 | 0.430 | 0.851 | -1.238 | 2.098 | 0.613 |
| Acrophase – Spring 2024 | 2.127 | 0.422 | 1.301 | 2.954 | <0.001 |
| **c. Sleep duration** | | | | | |
| | Estimate | SE | Lower CI | Upper CI | p |
| Intercept | 441.638 | 22.314 | 397.903 | 485.373 | <0.001 |
| Type of day (weekend) | 26.593 | 2.997 | 20.719 | 32.466 | <0.001 |
| Sex | -27.799 | 11.154 | -49.66 | -5.937 | 0.013 |
| Age | 1.321 | 0.562 | 0.219 | 2.422 | 0.019 |
| Winter 2021 | 9.264 | 5.938 | -2.375 | 20.903 | 0.119 |
| Spring 2024 | -14.678 | 13.466 | -41.072 | 11.715 | 0.276 |
| **Amplitude – Spring 2020** | **7.954** | **3.67** | **0.761** | **15.146** | **0.030** |
| Amplitude – Winter 2021 | 6.986 | 6.479 | -5.712 | 19.684 | 0.281 |
| **Amplitude – Spring 2024** | **9.055** | **2.454** | **4.245** | **13.866** | **<0.001** |
| Acrophase – Spring 2020 | -2.833 | 0.465 | -3.746 | -1.921 | <0.001 |
| Acrophase – Winter 2021 | 1.079 | 0.933 | -0.75 | 2.907 | 0.248 |
| Acrophase – Spring 2024 | -0.047 | 0.28 | -0.597 | 0.502 | 0.866 |

**Supplementary Table 3. Summary of the cosinor models testing lunar rhythmicity in Toba/Qom's sleep patterns.** The models include only one sinusoidal component with a period of 30 days. In each, one sleep outcome (sleep onset, offset, or duration) was included as the dependent variable, while night order in the lunar synodic cycle was the time variable and the cohort as the group. Additionally, we control for type of day (weekday/weekend), cohort, sex and age, including them as modifiers of the mesor. Winter 2016 cohort and weekdays are the reference levels. Significant amplitudes are shown in bold font. The intercept, amplitudes and the effects of sex, age and type of day are reported in hours for sleep onset and offset and in minutes for sleep duration. Acrophases are reported in radians. **a. Sleep onset.** Only the Fall 2023 cohort did not show a significant amplitude. Sleep onsets were represented as hours from midnight. **b. Sleep offset.** Both Fall 2023 and Spring 2023-Summer 2024 cohorts did not show a significant amplitude. Sleep offsets were represented as hours from midnight. **c. Sleep duration.** Both Fall 2023 and Spring 2023-Summer 2024 cohorts did not show a significant amplitude. Sleep duration was measured in minutes.

| **a. Sleep onset** | | | | | |
|---|---|---|---|---|---|
| | Estimate (h) | SE | Lower CI | Upper CI | p |
| Intercept | -1.921 | 0.202 | -2.317 | -1.525 | <0.001 |
| Type of day (weekend) | 0.183 | 0.039 | 0.106 | 0.260 | <0.001 |
| Sex | 0.556 | 0.167 | 0.229 | 0.882 | 0.001 |
| Age | 0.014 | 0.008 | -0.003 | 0.031 | 0.101 |
| Fall 2023 | 0.791 | 0.113 | 0.569 | 1.013 | <0.001 |
| Fall 2024 | 0.919 | 0.111 | 0.701 | 1.137 | <0.001 |
| Spring 2018 | 0.515 | 0.093 | 0.334 | 0.697 | <0.001 |
| Spring 2023 | 1.335 | 0.105 | 1.129 | 1.541 | <0.001 |
| Summer 2024 | 1.705 | 0.124 | 1.462 | 1.948 | <0.001 |
| **Amplitude – Winter 2016** | **0.176** | **0.069** | **0.042** | **0.311** | **0.010** |
| **Amplitude – Fall 2023** | **0.193** | **0.073** | **0.051** | **0.336** | **0.008** |
| **Amplitude – Fall 2024** | **0.160** | **0.054** | **0.054** | **0.265** | **0.003** |
| **Amplitude – Spring 2018** | **0.242** | **0.067** | **0.111** | **0.373** | **<0.001** |
| **Amplitude – Spring 2023** | **0.194** | **0.050** | **0.096** | **0.292** | **<0.001** |
| Amplitude – Summer 2024 | 0.170 | 0.103 | -0.032 | 0.372 | 0.100 |
| Acrophase – Winter 2016 | -0.324 | 0.408 | -1.123 | 0.475 | 0.427 |
| Acrophase – Fall 2023 | -1.763 | 0.424 | -2.593 | -0.932 | <0.001 |
| Acrophase – Fall 2024 | -2.921 | 0.303 | -3.515 | -2.326 | <0.001 |
| Acrophase – Spring 2018 | -1.535 | 0.262 | -2.048 | -1.022 | <0.001 |
| Acrophase – Spring 2023 | 1.621 | 0.252 | 1.126 | 2.116 | <0.001 |
| Acrophase – Summer 2024 | -2.693 | 0.631 | -3.930 | -1.457 | <0.001 |
| **b. Sleep offset** | | | | | |
| | Estimate (h) | SE | Lower CI | Upper CI | P |
| Intercept | 7.805 | 0.219 | 7.376 | 8.233 | <0.001 |
| Type of day (weekend) | 0.143 | 0.036 | 0.071 | 0.214 | <0.001 |
| Sex | 0.391 | 0.179 | 0.039 | 0.743 | 0.029 |
| Age | -0.035 | 0.009 | -0.053 | -0.016 | 0.000 |
| Fall 2023 | 0.474 | 0.111 | 0.256 | 0.692 | <0.001 |
| Fall 2024 | 0.502 | 0.111 | 0.284 | 0.719 | <0.001 |
| Spring 2018 | -0.041 | 0.087 | -0.213 | 0.130 | 0.635 |
| Spring 2023 | 0.033 | 0.105 | -0.172 | 0.238 | 0.754 |
| Summer 2024 | 0.586 | 0.121 | 0.348 | 0.824 | <0.001 |
| **Amplitude – Winter 2016** | **0.170** | **0.068** | **0.038** | **0.303** | **0.012** |
| Amplitude – Fall 2023 | 0.080 | 0.071 | -0.059 | 0.219 | 0.257 |
| **Amplitude – Fall 2024** | **0.226** | **0.047** | **0.134** | **0.318** | **<0.001** |

| | | | | | |
|---|---|---|---|---|---|
| **Amplitude – Spring 2018** | **0.150** | **0.060** | **0.032** | **0.267** | **0.012** |
| Amplitude – Spring 2023 | 0.077 | 0.049 | -0.020 | 0.174 | 0.118 |
| Amplitude – Summer 2024 | 0.127 | 0.089 | -0.046 | 0.301 | 0.151 |
| Acrophase – Winter 2016 | -1.645 | 0.387 | -2.403 | -0.886 | <0.001 |
| Acrophase – Fall 2023 | -2.173 | 0.962 | -4.057 | -0.288 | 0.024 |
| Acrophase – Fall 2024 | 1.198 | 0.227 | 0.753 | 1.643 | <0.001 |
| Acrophase – Spring 2018 | 0.645 | 0.429 | -0.197 | 1.487 | 0.133 |
| Acrophase – Spring 2023 | 1.886 | 0.590 | 0.729 | 3.042 | 0.001 |
| Acrophase – Summer 2024 | -2.077 | 0.848 | -3.738 | -0.415 | 0.014 |

**c. Sleep duration**

| | Estimate (min) | SE | Lower CI | Upper CI | p |
|---|---|---|---|---|---|
| Intercept | 583.297 | 9.979 | 563.738 | 602.855 | <0.001 |
| Type of day (weekend) | -2.440 | 2.743 | -7.816 | 2.937 | 0.374 |
| Sex | -7.552 | 7.648 | -22.542 | 7.438 | 0.323 |
| Age | -3.002 | 0.390 | -3.765 | -2.238 | <0.001 |
| Fall 2023 | -17.839 | 6.967 | -31.494 | -4.185 | 0.010 |
| Fall 2024 | -22.449 | 6.495 | -35.179 | -9.718 | 0.001 |
| Spring 2018 | -34.041 | 6.051 | -45.901 | -22.180 | <0.001 |
| Spring 2023 | -76.433 | 6.199 | -88.583 | -64.284 | <0.001 |
| Summer 2024 | -65.041 | 7.737 | -80.206 | -49.876 | <0.001 |
| **Amplitude – Winter 2016** | **12.682** | **4.859** | **3.157** | **22.206** | **0.009** |
| Amplitude – Fall 2023 | 8.861 | 4.913 | -0.769 | 18.491 | 0.071 |
| **Amplitude – Fall 2024** | **20.363** | **3.661** | **13.188** | **27.538** | **<0.001** |
| **Amplitude – Spring 2018** | **21.072** | **4.626** | **12.005** | **30.139** | **<0.001** |
| **Amplitude – Spring 2023** | **7.302** | **3.425** | **0.588** | **14.015** | **0.033** |
| Amplitude – Summer 2024 | 3.979 | 7.763 | -11.235 | 19.193 | 0.608 |
| Acrophase – Winter 2016 | -2.604 | 0.380 | -3.350 | -1.859 | <0.001 |
| Acrophase – Fall 2023 | 1.820 | 0.642 | 0.560 | 3.079 | 0.005 |
| Acrophase – Fall 2024 | 0.796 | 0.165 | 0.473 | 1.119 | <0.001 |
| Acrophase – Spring 2018 | 1.174 | 0.202 | 0.778 | 1.570 | <0.001 |
| Acrophase – Spring 2023 | -1.641 | 0.459 | -2.540 | -0.741 | <0.001 |
| Acrophase – Spring 2024 | -0.418 | 1.679 | -3.709 | 2.872 | 0.803 |

**Supplementary Table 4. Summary of the cosinor models testing lunar rhythmicity in the activity onset of captive titi monkeys.** The models include only one sinusoidal component with a period of 30 days. Activity onset was included as the dependent variable, night in the lunar synodic cycle as the time variable and cohort as the group. Additionally, we control for cohort, including them as modifiers of the mesor. Summer 2023 is the reference. Significant amplitudes are shown in bold font. The intercept and amplitudes are reported in hours, and acrophases are reported in radians.

| | Estimate | SE | Lower CI | Upper CI | p |
|---|---|---|---|---|---|
| Intercept | 5.389 | 0.063 | 5.265 | 5.514 | <0.001 |
| Spring 2024 | 0.245 | 0.090 | 0.069 | 0.421 | 0.006 |
| **Amplitude – Summer 2023** | **0.120** | **0.027** | **0.067** | **0.172** | **<0.001** |
| **Amplitude – Spring 2024** | **0.088** | **0.031** | **0.027** | **0.150** | **0.005** |
| Acrophase – Summer 2023 | -1.347 | 0.241 | -1.819 | -0.874 | <0.001 |
| Acrophase – Spring 2024 | 2.217 | 0.370 | 1.493 | 2.942 | <0.001 |

### Supplementary Figures

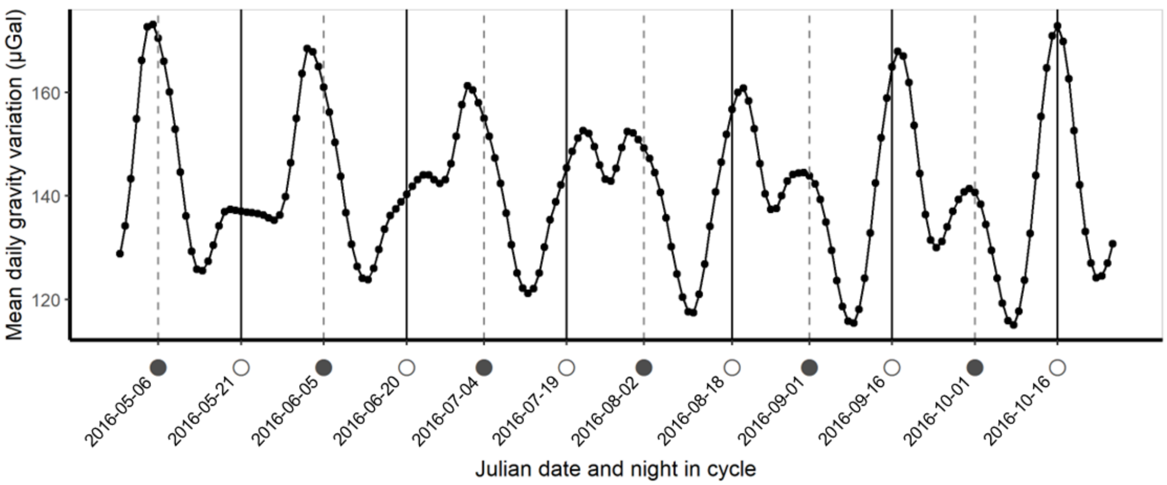


**Supplementary Figure 1. Changes in mean daily gravitational variation caused by the Moon and the Sun over several lunar synodic cycles.** The figure displays the simulated values for the mean gravitational variation at Ingeniero Juárez, Formosa, Argentina, between April 29 and October 26 of 2016. New moons are represented as dark circles and full moons as white circles. Peaks of gravitational variation happen around new and full moons, but their magnitude varies every synodic month. For example, during May, there was an absolute maximum around the new moon and a lower local maximum around the full moon; the exact opposite happened in October.

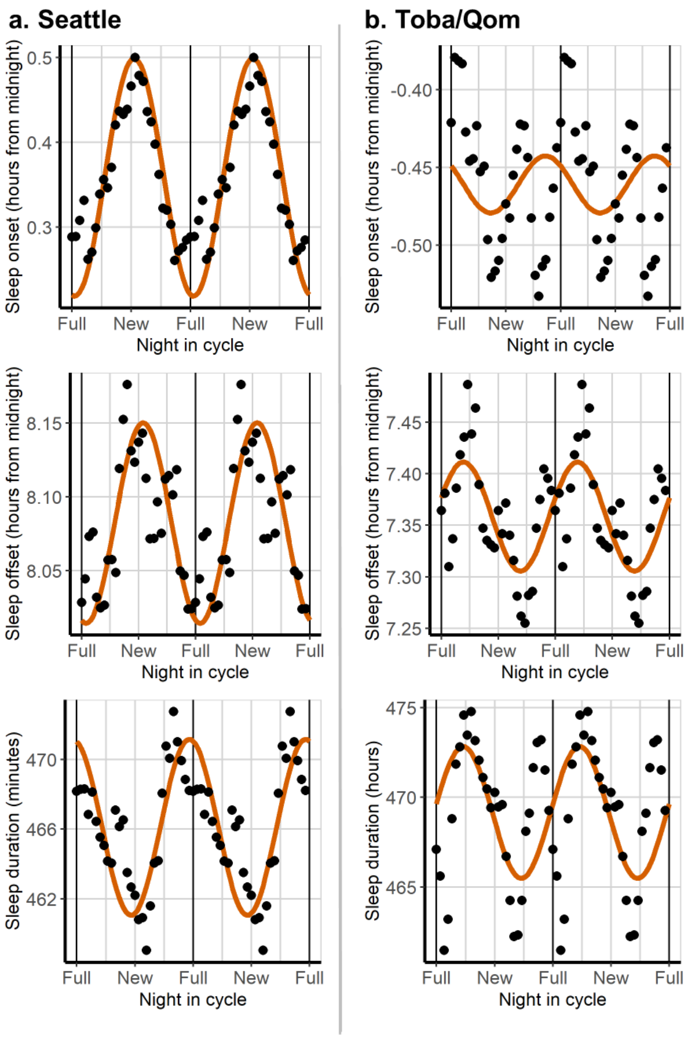


**Supplementary Figure 2. Lunar regulation of offset and duration in Seattle and in Toba/Qom communities.** For each sleep variable in each population, we adjusted a cosinor model with a 30-day period to study lunar rhythmicity. We included participant's id as a random factor. Orange lines show the prediction of the cosinor models. Dots represent the mean value of sleep parameters by night in cycle. **a. Seattle.** Sleep onset presents lunar rhythmicity, that is, the amplitude of the adjusted model was significantly different from zero (a= 0.13 h, 95% CI = 0.45-0.22, p=0.003), showing later bedtimes around the new moon and earlier around the full moon. Sleep offset did not show a significant amplitude (a= 0.07 h, 95% CI = -0.02-0.15, p=0.124). Sleep duration presents lunar rhythmicity, that is, the amplitude of the adjusted model was significantly different from zero (sleep duration: a=4.38 min, 95% CI = 0.19-7.96 p=0.017). However, the phase was opposite to what was previously reported, showing later bedtimes and shorter sleep duration around the new moon instead of the full moon. Additionally, the amplitudes found here are smaller than previously reported. **b. Toba/Qom communities.** Sleep onset did not show rhythmicity in Toba/Qom communities (a=0.024 h, 95% CI = -0.026-0.074, p= 0.346). Additionally, neither sleep offset nor sleep duration showed lunar rhythmicity in Toba/Qom communities (sleep offset: a=0.057 h, 95% CI = 0.007-0.107, p=0.026; and sleep duration: a= 4.62 min, 95% CI= 1.16-8.09, p= 0.010).

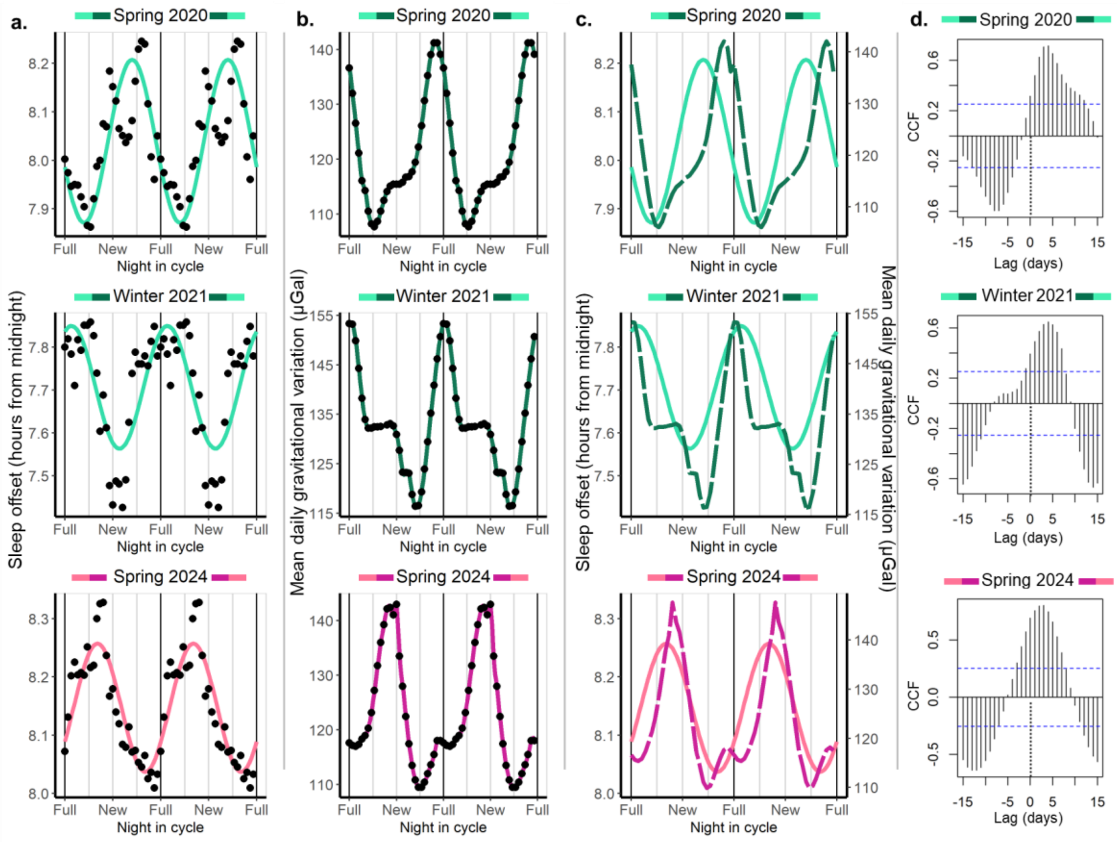


**Supplementary Figure 3. Sleep offset in Seattle shows lunar rhythmicity, its phase depends on the cohort, and it is highly correlated with the mean daily gravitational variation caused by the Moon and the Sun.** The colour assigned to each cohort is based on the shape of the mean daily gravitational variation and its phase relationship with the synodic lunar month. Green represents a clear maximum peak associated with the full moon, while pink indicates a clear maximum peak associated with the new moon. **a. Sleep offset as a function of night in the synodic Lunar cycle.** Phases differ between cohorts, with the Spring of 2020 and the Winter of 2021 presenting later sleep offsets around/before the full Moon, and the Spring of 2024 presenting later wake-up times before the new Moon. According to the adjusted model, the Spring of 2020 was the only cohort with an amplitude that differed from zero. Data are double-plotted. Dots represent the mean value by night in cycle to which a 7-day moving average was applied. The colored line represents the prediction of the fitted mixed cosinor model describing the relationship between night in cycle and sleep offset, where the cohort was included as a predictor. Additionally, we controlled by type of day (weekend or weekday), sex and age, and we included the id of the participants as a random factor. **b. Mean daily gravitational variation (including Moon and Sun effects).** The absolute maximums of the mean daily gravitational variation differ between cohorts, happening around the full Moon for the Spring of 2020 and the Winter of 2021, and around the new Moon for the Spring of 2024. Dots represent average values by night in cycle for the same dates that were included in (a) The colored lines are for visualization; they smoothly connect consecutive dots. **c. Sleep offset and mean daily gravitational variation show similar patterns.** Colored full lines represent the prediction of the model for sleep offset, and colored dotted lines represent the mean daily gravitational variation. The peaks of sleep precede the peaks of mean daily gravitational variation, or they are highly coincident with them. **d. Cross-correlations between sleep offset and the mean daily gravitational variation at the cohort level.** Plots show the cross-correlations for each cohort using the average values by night in cycle, that is, the cross-correlations between the dots represented in (a) and (b). Results are consistent between cohorts, showing a positive correlation between mean daily gravitational variation and sleep offset around or before lag zero.

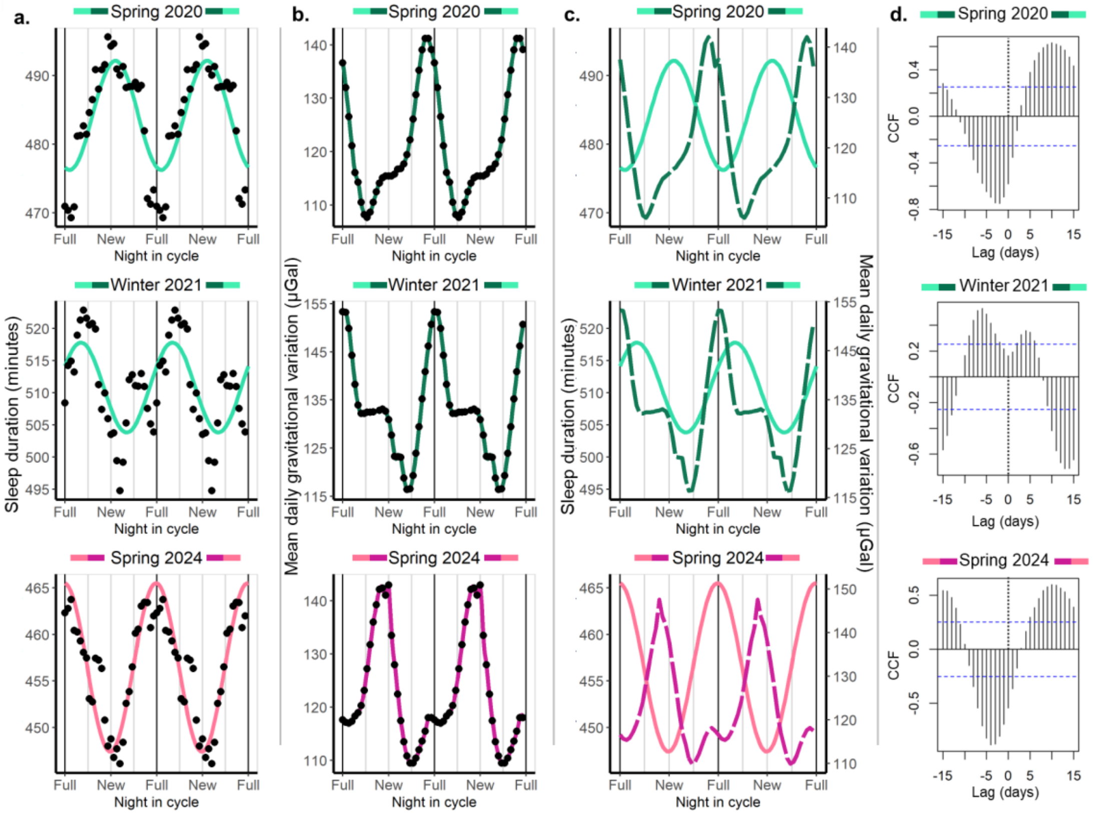


**Supplementary Figure 4. Sleep duration in Seattle shows lunar rhythmicity, its phase depends on the cohort, and it is highly correlated with the mean daily gravitational variation caused by the Moon and the Sun.** The colour assigned to each cohort is based on the shape of the mean daily gravitational variation and its phase relationship with the synodic lunar month. Green represents a clear maximum peak associated with the full moon while pink indicates a clear maximum peak associated with the new moon. **a. Sleep duration as a function of night in the synodic Lunar cycle.** According to the adjusted model, the Winter of 2021 did not show an amplitude that differs from zero. For the other two cohorts, phases differ, with the Spring of 2020 presenting shorter sleep durations close to the full Moon and the Spring of 2024 presenting shorter sleep durations around the new Moon. Data are double-plotted. Dots represent the mean value by night in cycle to which a 7-day moving average was applied. The colored line represents the prediction of the fitted mixed cosinor model describing the relationship between night in cycle and sleep duration, where the cohort was included as a predictor. Additionally, we controlled by type of day (weekend or weekday), sex and age, and we included the id of the participants as a random factor. **b. Mean daily gravitational variation (including Moon and Sun effects).** The absolute maximums of the mean daily gravitational variation differ between cohorts, happening around the full Moon for the Spring of 2020 and the Winter of 2021, and around the new Moon for the Spring of 2024. Dots represent average values by night in cycle for the same dates that were included in (b). The colored lines are for visualization, they smoothly connect consecutive dots. **c. Sleep duration and the mean daily gravitational variation show opposite patterns.** Colored full lines represent the prediction of the model for sleep duration, and colored dotted lines represent the mean daily gravitational variation. The peaks of sleep duration coincide with the troughs of mean daily gravitational variation for the Spring of 2020 and 2024. **d. Cross-correlations between sleep duration and mean daily gravitational variation at the cohort level.** Plots show the cross-correlations for each cohort using the average values by night in cycle, that is, the cross-correlations between the dots represented in (a) and (b). Results are consistent between Spring 2020 and 2024, showing a negative correlation between mean daily gravitational variation and sleep duration close to lag zero.

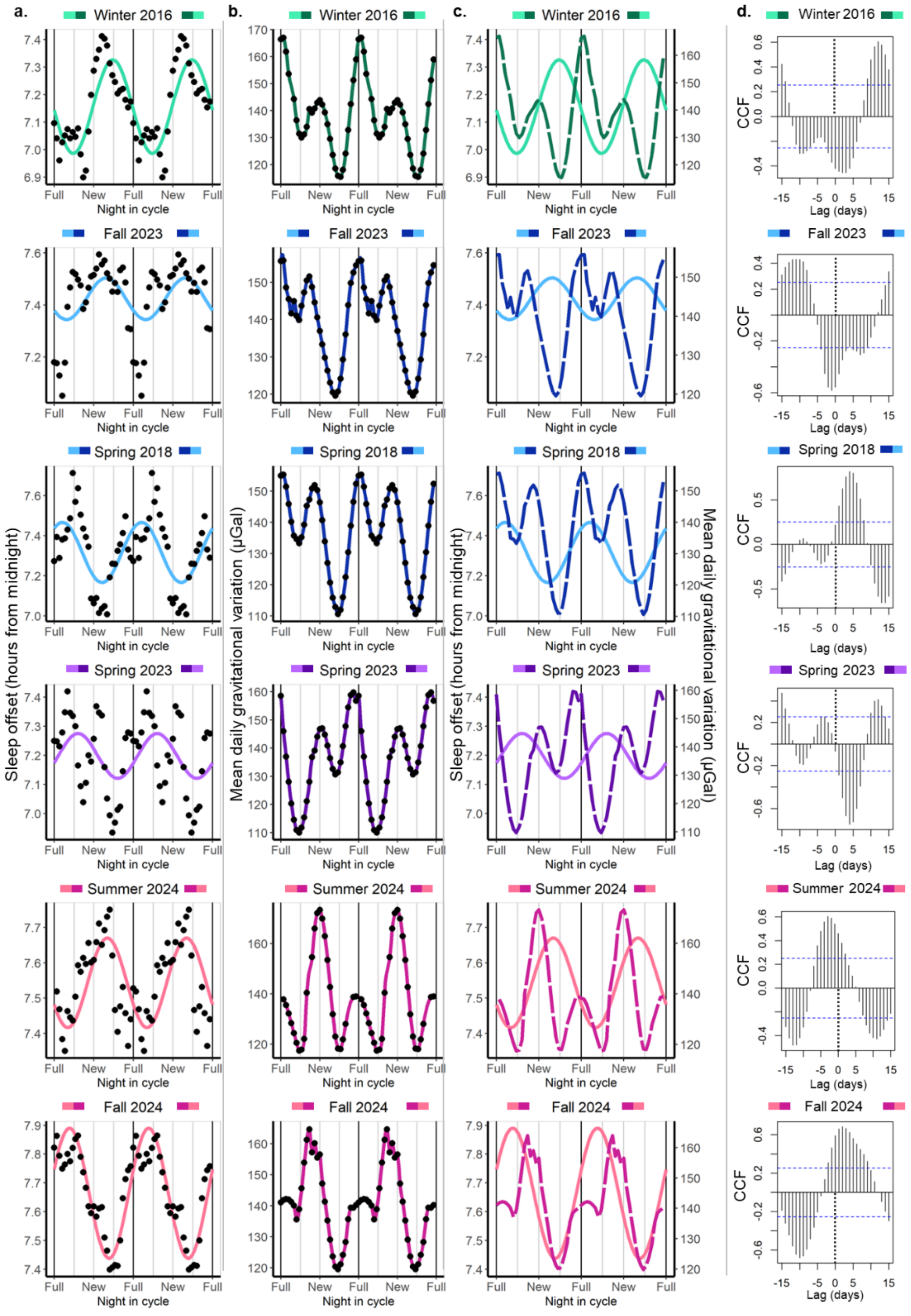


**Supplementary Figure 5. Sleep offset in Toba/Qom communities shows lunar rhythmicity, its phase depends on the cohort, and it is correlated with the mean daily gravitational variation caused by the Moon and the Sun.** The colour assigned to each cohort is based on the shape of the mean daily gravitational variation and its phase relationship with the synodic lunar month. Green represents a clear maximum peak associated with the full moon, while pink indicates a clear maximum peak associated with the new moon. When peaks around the full and new moon were similar, we used blue to identify cohorts with an absolute minimum before the full moon and purple to identify cohorts with an absolute minimum before the new moon. **a. Sleep offset as a function of night in the synodic Lunar cycle.** According to the adjusted model, the cohorts whose amplitude differed from zero were Spring 2012, Spring 2018 and Fall 2024. Dots represent the mean value by night in cycle to which a 7-day moving average was applied. The coloured line represents the prediction of the fitted mixed cosinor model describing the relationship between night in cycle and sleep offset, where the cohort was included as a predictor. Additionally, we controlled by type of day (weekend or weekday), sex and age, and we included the id of the participants as a random factor. **b. Mean daily gravitational variation (including Moon and Sun effects).** The patterns of the mean daily gravitational variation differ between cohorts, with at least 4 different patterns being present. Plots were organized according to these patterns. Dots represent average values by night in cycle for the same dates that were included in (a). The coloured lines are for visualization, and it is a spline-based smoothing that connects consecutive dots. **c. Sleep offset and mean daily gravitational variation.** Coloured full lines represent the prediction of the model for sleep offset and coloured dotted lines represent the mean daily gravitational variation. **d. Cross-correlations between sleep offset and the mean daily gravitational variation at the cohort level.** Plots showed the cross-correlations for each cohort using the average values by night in cycle, that is, the cross-correlations between the dots represented in (a) and (b).

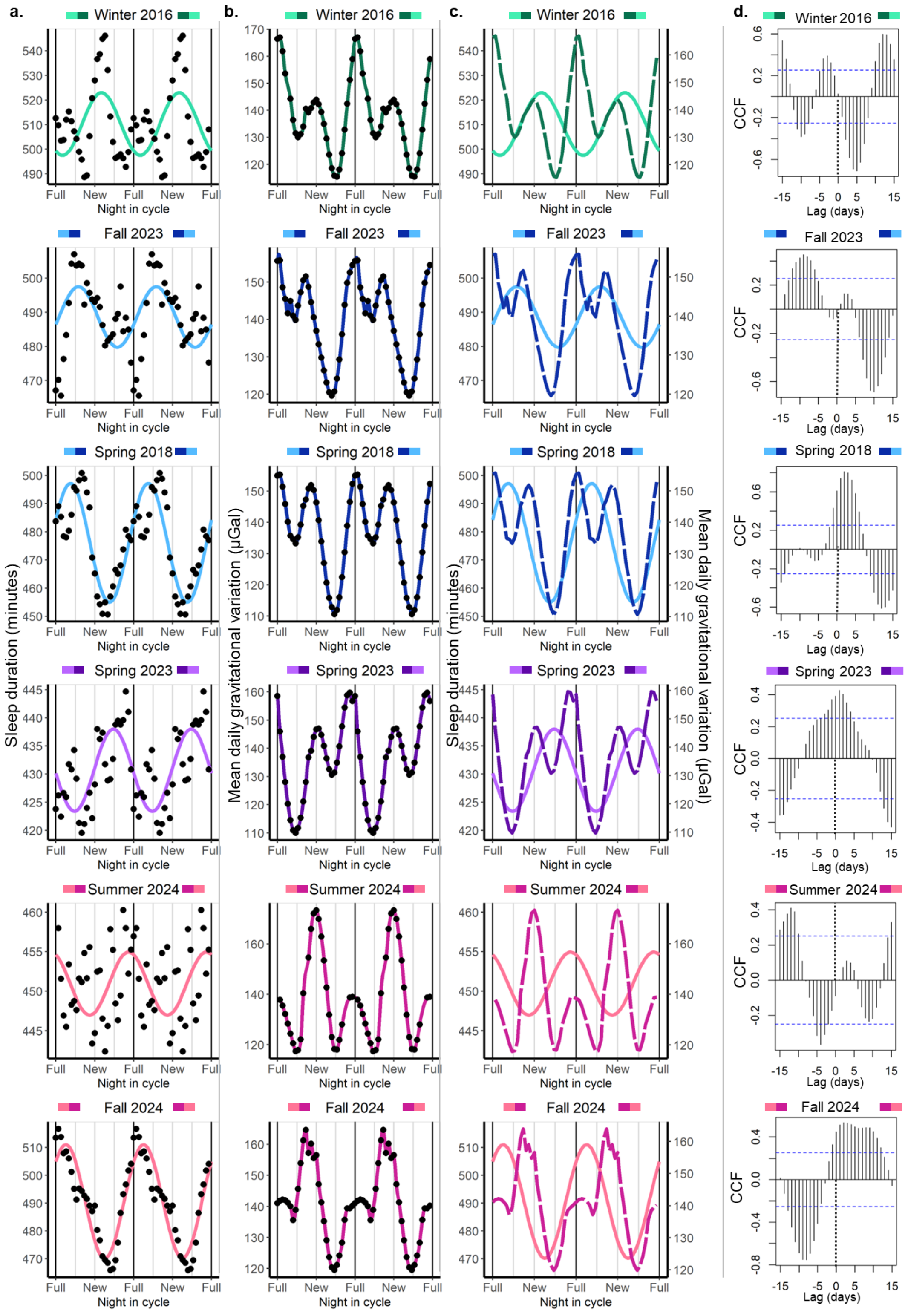


**Supplementary Figure 6. Sleep duration in Toba/Qom communities shows lunar rhythmicity, its phase depends on the cohort, and it is correlated with the mean daily gravitational variation caused by the Moon and the Sun.** The colour assigned to each cohort is based on the shape of the mean daily gravitational variation and its phase relationship with the synodic lunar month. Green represents a clear maximum peak associated with the full moon, while pink indicates a clear maximum peak associated with the new moon. When peaks around the full and new moon were similar, we used blue to identify cohorts with an absolute minimum before the full moon and purple to identify cohorts with an absolute minimum before the new moon. **a. Sleep duration as a function of night in the synodic Lunar cycle.** According to the adjusted model, the cohorts whose amplitude differ from zero were Winter 2016, Spring 2018, Spring 2023 and Fall 2024. Data are double-plotted. Dots represent the mean value by night in cycle to which a 7-day moving average was applied. The coloured line represents the prediction of the fitted mixed cosinor model describing the relationship between night in cycle and sleep duration, where the cohort was included as a predictor. Additionally, we controlled by type of day (weekend or weekday) sex and age, and we included the id of the participants as a random factor. **b. Mean daily gravitational variation (including Moon and Sun effects).** Dots represent average values by night in cycle for the same dates that were included in (a) The coloured lines are for visualisation, they smoothly connect consecutive dots. **c. Sleep duration and mean daily gravitational variation show similar patterns.** Coloured full lines represent the prediction of the model for sleep duration, and coloured dotted lines represent the mean daily gravitational variation. **d. Cross-correlations between sleep duration and mean daily gravitational variation at the cohort level.** Plots showed the cross-correlations for each cohort using the average values by night in cycle, that is, the cross-correlations between the dots represented in (a) and (b).

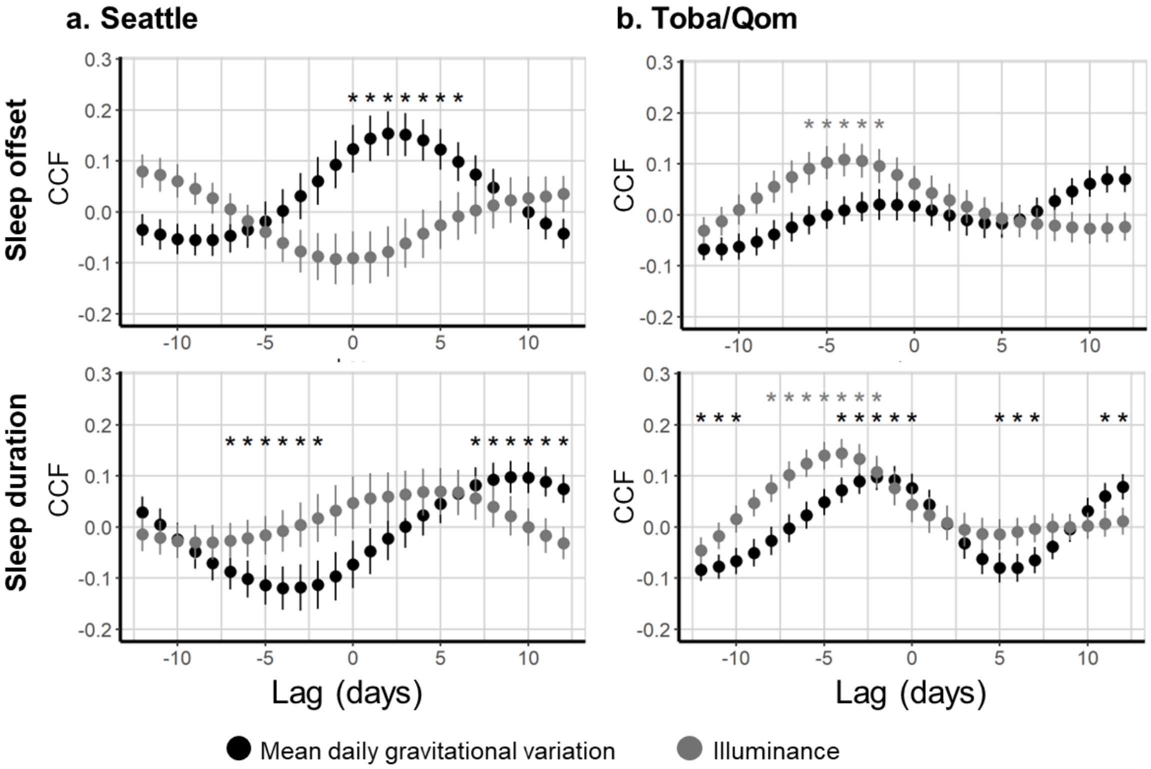


**Supplementary Figure 7. Cross-correlations of the mean daily gravitational variation and Moon's illuminance with sleep offset and duration in Seattle and Toba/Qom participants.** We calculated the cross-correlation at the individual level between different sleep outcomes (sleep onset, offset and duration) and the mean daily gravitational variation, on the one hand, and between these sleep outcomes and the Moon's illuminance, on the other hand. The plots showed the mean CCF for each lag and its standard error. Asterisks show whether mean CCFs by lag differ from zero after controlling for multiple comparisons. **a. Seattle.** Cross-correlations between sleep offset and the mean daily gravitational variation follow a similar pattern to those shown for sleep onset: later sleep offsets precede higher gravitational variations. No CCF was statistically different from zero for sleep offset and moon's illuminance. CCFs for sleep duration and the mean daily gravitational variation follow the opposite pattern to that shown for sleep onset, with a shift to the left (e.g. shorter sleep durations happening after higher mean daily gravitational variations). CCFs for sleep duration and moon's illuminance were not significantly different from zero. **b. Toba/Qom.** While no CCF for sleep offset and the mean daily gravitational variation was statistically significant, positive correlations were found for the Moon's illuminance for lags between -6 and -2. On the one hand, cross-correlations for sleep duration and the mean daily gravitational pull follow the opposite pattern to those shown for sleep onset. On the other hand, the association between sleep duration and the Moon's illuminance, the observed pattern was similar to the one observed for sleep offset and Moon's illuminance. Positive CCFs were found for lags -4-0 and 12-14 for mean daily gravitational variation, and for lags -8 to -2 for moon's illuminance and sleep duration.
